# Motion of the Cochlear Reticular Lamina Varies Radially Across Outer-Hair-Cell Rows

**DOI:** 10.1101/2022.03.01.482580

**Authors:** Nam Hyun Cho, Sunil Puria

## Abstract

The basilar membrane (BM) is connected to the reticular lamina (RL) through three rows of Y-shaped structures consisting of an outer hair cell (OHC) and a Deiters’ cell (DC) with a phalangeal process (PhP) that forms part of the RL mosaic surface. Morphological differences in the anatomy of the Y-shaped structures across the three OHC rows suggest differences in motion across the rows. Here we report OoC transverse motions measured across several radial locations for the gerbil basal region corresponding to ~45 kHz. Cross-sectional imaging and vibrometry measurements were made using a high-resolution (2.23 um axially in water) spectral-domain optical-coherence-tomography (SD-OCT) system. The stimuli were pure tones (2–63 kHz) at ear-canal sound pressure levels (SPLs) of 30–95 dB SPL in anesthetized gerbils (N=9) with healthy cochleae. We report displacements at the RL regions of OHC rows 1–3 (RL_1–3_), at the OHC-DC junctions of OHC rows 1–3 (OHC-DC-junction_1–3_), and at the arcuate zone, arcuate-pectinate junction, and pectinate zone of the BM (BM_AZ_, BM_APJ_, and BM_PZ_, respectively). The in vivo BM displacements showed classic compressive nonlinearity and traveling-wave delays. The RL gain was similar to the BM gain at low frequencies (<20 kHz), but increased with frequency. Near the best frequency (BF), the RL gain was greater than the high-level BM gain by 40 ±5 dB (mean±std), and had greater compressive nonlinearity. RL motion varied radially, and the RL_3_ gain was significantly greater than that of RL_1_ by 10 ±1 dB (p<0.001). In contrast, the OHC-DC-junction gain varied little radially across OHCs. At low frequencies the OHC-DC-junction gain was constant across SPLs, and 14 ± 3 dB greater than the BM gain. As the frequency increased, the OHC-DC-junction gain decreased to a level similar to the BM gain at BF. The RL_2, 3_ phase was advanced by 0.25–0.375 cycles relative to the BM phase at low frequencies, but the RL_2, 3_ phase lead decreased as the frequency increased, became similar to the BM phase at BF, and lagged behind the BM phase by 0.25–0.5 cycles above BF. The OHC-DC-junction phases were mostly similar to the BM phase at low frequencies, but became delayed relative to the BM as the frequency increased, typically by 0.25–0.5 cycles near BF and by up to 1 cycle above BF. Our results show the most detailed picture of motion around the three OHC rows yet published, indicating that RL motion varied radially. Surprisingly, there was little motion difference across the three OHC rows in the OHC-DC-junction region, indicating that the tops of the DCs move in unison. Our data show a rich array of OoC amplitude and phase variations that are not explained by current theories.

## Introduction

The great sensitivity and frequency selectivity of mammalian hearing originates in the mechanical properties of the cochlea. Cochlear mechanical motions in response to sound are amplified using metabolic energy. The motor element of this cochlear amplification is the outer hair cell (OHC), that can change its length at sound frequencies. Each basally-tilted OHC, along with its attached Deiters’ cell (DC) and the long apically-tilted DC phalangeal process (PhP), form a Y-shaped building block of the complex mechanical skeleton (Soons et al., 2015) occupying the space between the reticular lamina (RL) and the basilar membrane (BM) within the organ of Corti (OoC). How the OHCs, acting through this skeleton of Y-shaped structures, work to achieve cochlear amplification is not fully understood.

In the classic view of the cochlea, BM motion was considered the most important, and the motions of the rest of the OoC were thought to follow the BM motion (reviews: Robles and Ruggero, 2001; Guinan et al., 2012). However, recently developed optical methods have allowed motion measurements within the OoC of live, sensitive animals, and have revealed a much different picture. In response to sound, motions of other structures within the OoC are much larger than BM motion, and have motion responses that are amplified over a larger frequency range than the BM (e.g., Chen et al., 2011; Ramamoorthy et al., 2014; Lee et al., 2015; 2016; Ren et al., 2016; Fallah et al., 2019).

Another recent development is the realization that the phasing of the OHC motion required to amplify BM motion does not arise from a resonance in the tectorial membrane (TM; Dong and Olson 2013; Lee et al., 2016; Guinan, 2020), but is instead arises from an unexpected phase of the radial motion of the RL. The RL radial-motion phase measured by Lee et al. (2016) was almost opposite to the phase expected from the OoC rotation resulting from BM motion. How such an RL phase might be produced is unknown. The RL surface is a *mosaic* comprised of the cuticular plates of the basally tilted OHCs and apically tilted PhPs, all held together by adhesion molecules (Aijaz et al., 2006; Nunes et al., 2006), and as such it is likely to bend and/or stretch. It seems possible that skeleton of Y-shaped structures below the RL mosaic may play a role in creating the required RL radial-motion phase. The longitudinal and radial angles of the PhP branches of the Y-shaped structures vary across the three rows of OHCs (Soons et al., 2015), and this might cause the motions to be different across the three OHC rows, but this has yet to be experimentally or computationally demonstrated.

Most past work that measured motion in the OoC has concentrated on the motion profile along the direction corresponding to the OHC length change due to electromotility, which is the direction in which the OHC force acting against the BM is expected to produce BM cochlear amplification (e.g., Dewey et al, 2021). Here we examine OoC motion at various depths along this direction, and also at different radial locations, namely at each of the three rows of OHCs. The results should help in understanding how the OoC works, and may shed light on the mechanisms by which the RL radial motion could achieve the correct phase for cochlear amplification.

## Results

The location relative to the anatomy in the gerbil hook region where vibration measurements were made is shown in Figure 1a–c. At the wavelength of our optical coherence tomography (OCT) system, there were reflectivity peaks suitable for making measurements at the RL, at the bottom of each OHC at its junction with the DC (OHC-DC-junction), and at the BM (Fig. 1d–g). At the RL and OHC-DC-junction points, it was often possible to see separate areas of high reflectivity that were aligned with each of the three OHC rows. The second-row area of high reflectivity was sometimes best seen at a slightly different longitudinal location, thus requiring a slight longitudinal offset (~15 μm) of the row 2 measurements relative to those of rows 1 and 3.

**Fig. 1.**
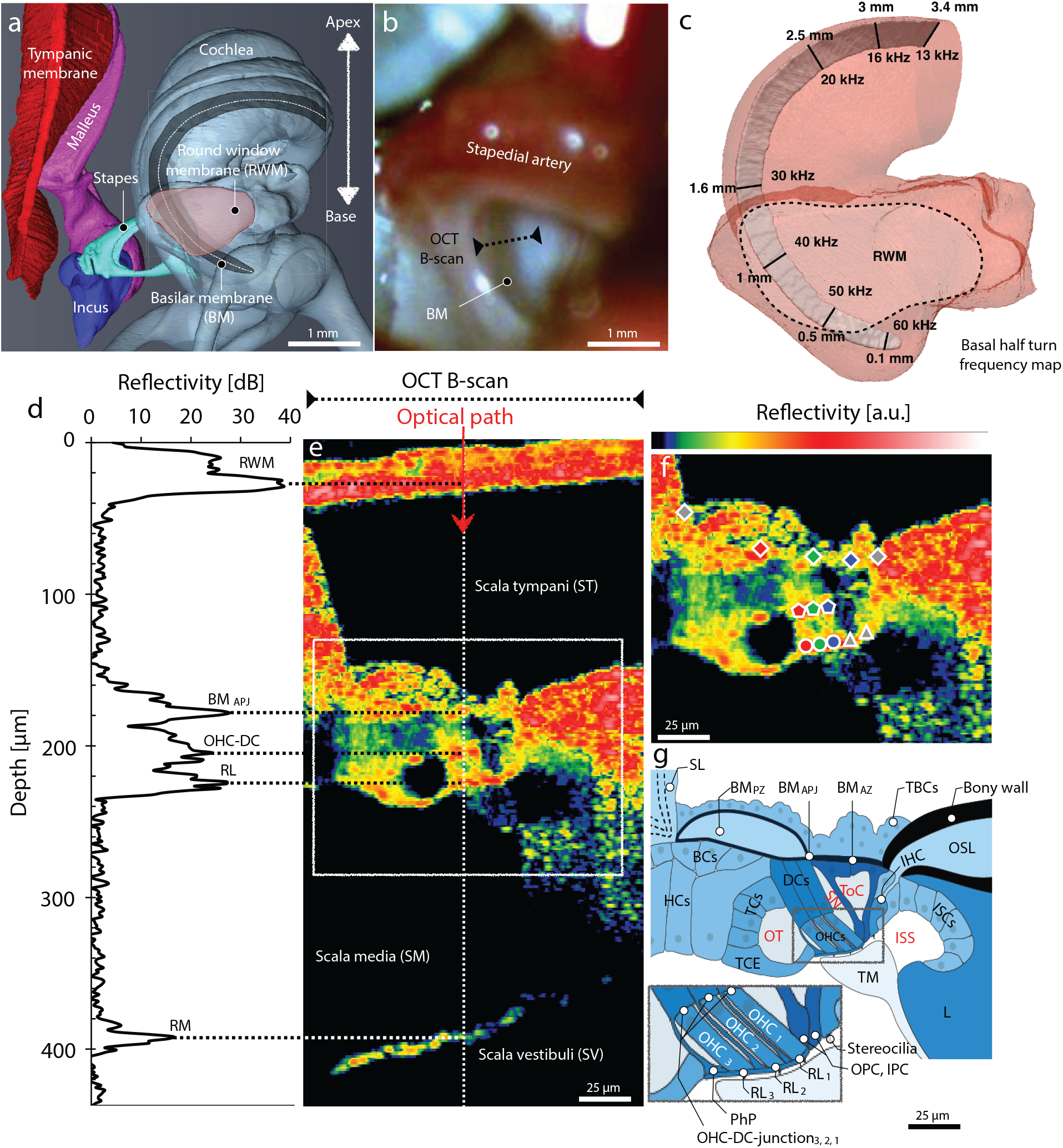
The anatomy of the gerbil ear and in vivo cochlea imaging. **a** A micro-computed-tomography (*μ*CT)–based 3D reconstruction shows the main structures of the gerbil middle ear (i.e., tympanic membrane, malleus, incus, and stapes), and the cochlea (base to apex) with the round-window membrane (RWM; pink region) and internal basilar membrane (BM; dark-gray region) indicated. **b** A photograph (specimen G504) shows the RWM region through which 2D “B-scan” cross sections of the BM in the basal “hook region” of the cochlea can be imaged using optical coherence tomography (OCT; black dotted line). **c** The frequency map of the BM “basal half turn” (from panel **a**) is shown as a function of distance from the basal tip. **d** An example of the depth profile (backscattered-light reflectivity) of a single 1D “A-scan” (white dotted line in **e**) shows several peaks corresponding to: the RWM; the junction (BM_APJ_) of the BM arcuate zone (BM_AZ_) and pectinate zone (BM_PZ_); the junction (OHC-DC) of an outer hair cell (OHC) and Deiters’ cell (DC); the reticular lamina (RL); and Reissner’s membrane (RM). **e** An in vivo OCT B-scan image (G612), as measured through the RWM. **f** Enlarged view of **e** detailing the organ of Corti (OoC) structure and marking the following OCT-vibrometry measurement points (left to right and top to bottom): outer BM edge, BM_PZ_, BM_APJ_, BM_AZ_, inner BM edge (gray/red/green/blue/gray diamonds); OHC-DC-junction_3,2,1_ (red/green/blue pentagons); RL_3,2,1_ (red/green/blue circles); and outer and inner pillar cells (OPC and IPC; gray triangles). **g** A labeled cross-sectional drawing of a representative OoC structure. The inset details the three rows of OHCs where they meet the RL at their apical ends, as well as the OPC, IPC, and inner-hair-cell (IHC) stereocilia. **Other abbreviations:** SL: spiral ligament; TBCs: tympanic border cells; BCs: Boettcher cells; OSL: osseous spiral lamina; HCs: Hensen’s cells; TCs: tectal cells; TCE: tectal-cell extension; ISCs: inner-sulcus cells; TM: tectorial membrane; L: limbus; OT: outer tunnel; SN: Space of Nuel; ToC: tunnel of Corti; ISS: inner spiral sulcus; PhP: phalangeal process.

### Motion measurements and gains

An example of motion displacement measurements from RL_3_ (the RL point at the top of OHC_3_, the third OHC row), along with the corresponding ear-canal sound pressure measurements, are shown in Figure 2. Also shown is the ratio (motion/sound-pressure), which we refer to as the motion “gain.” In all later figures, the motion displacement measurements are expressed in terms of their gain, which includes both a magnitude and a phase.

**Fig. 2.**
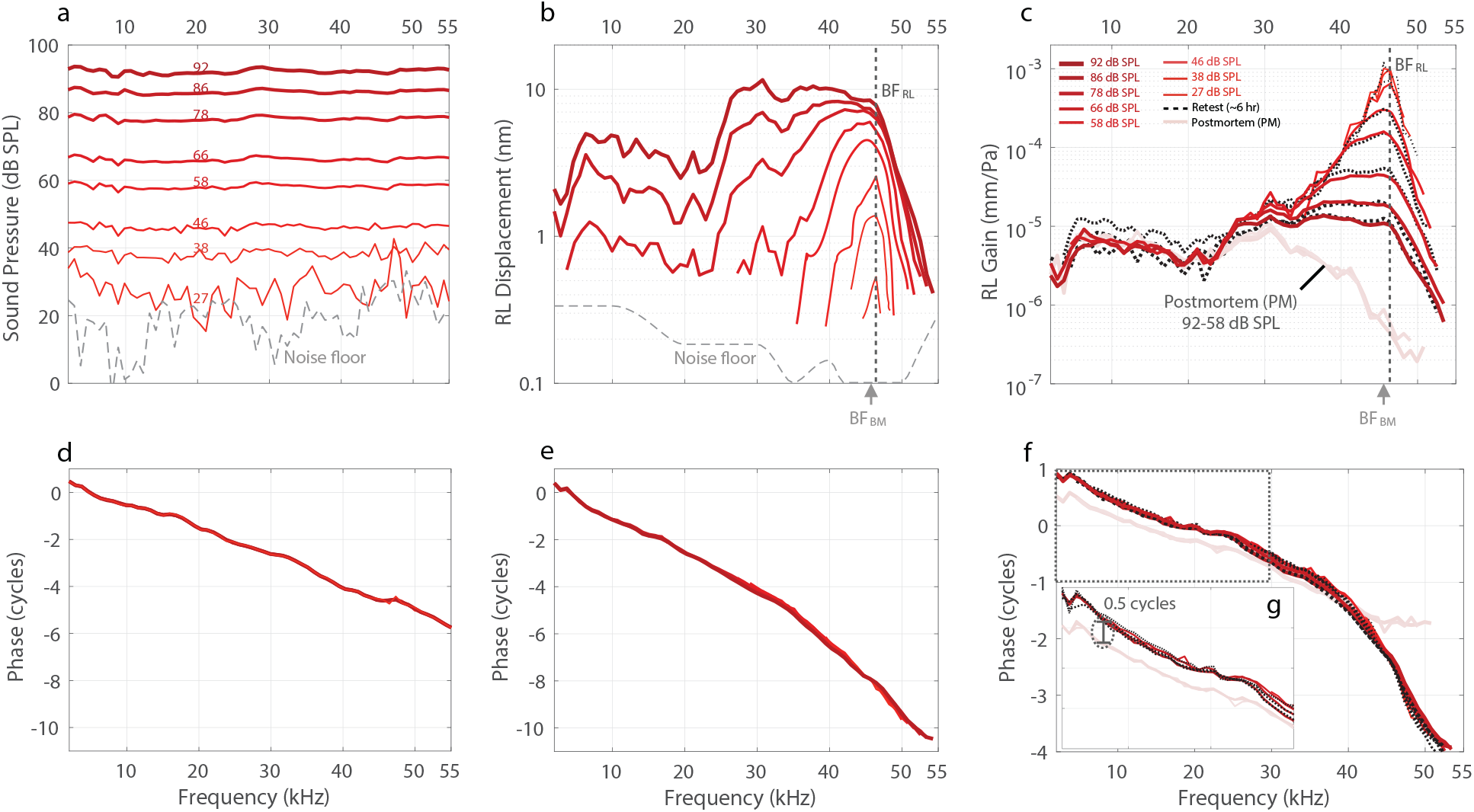
In vivo RL_3_ vibrometry measurements (see Fig. 1g inset) at different stimulus levels in the hook region of the gerbil cochlea (G637). **a** The ear-canal sound pressure levels of pure tones used to stimulate the specimen, ranging from 27 to 92 dB SPL (line thickness and darkness increase with level), as well as a noise-floor reading (gray dashed line). **b** RL_3_ displacements as functions of frequency, corresponding to the different stimulus levels in panel **a**. Displacement measurements were rejected at frequencies where their magnitude was less than 6 dB above the noise floor (gray dashed line). The best frequency (BF) for the RL_3_ location (BF_RL_) is 46.4 kHz (gray dotted vertical line), and for the BM_APJ_ location (BF_BM_) is 45.6 kHz (gray arrow below the panel). **c** The in vivo (dark red) and postmortem (PM; faded red) RL_3_ displacements are normalized by the stimulus sound pressure to produce the RL Gain (in units of mm/Pa). The black dotted lines indicate repeated in vivo measurements made ~6 hours later. **d–f** The respective phase responses of the sound pressure, RL_3_ displacement, and RL_3_ gain. **g** An enlarged view of the RL_3_ gain from the black-dotted region of panel **f**. s

Figure 2 shows that as the sound level went up, the RL_3_ motion grew compressively at frequencies near the best frequency (BF), but at low frequencies (<20 kHz) it grew close to linearly (i.e., the gains stayed the same, e.g., Figs. 2c, 3d, 4a). At BF, the RL motion had more than 60 dB of gain relative to the motion in the dead animal (Fig. 2c). This extraordinarily high gain demonstrates that this was a sensitive preparation. Repeated measurements made at the same location indicate that the in vivo specimen was stable even after 6 hours (Fig. 2c, dotted black and solid red lines). There was little variation in phase across sound levels (the red lines all overlap), but the phase did change from living to dead (Fig. 2f, g). At low frequencies there was almost a reversal of the RL_3_ motion phase.

**Fig. 3.**
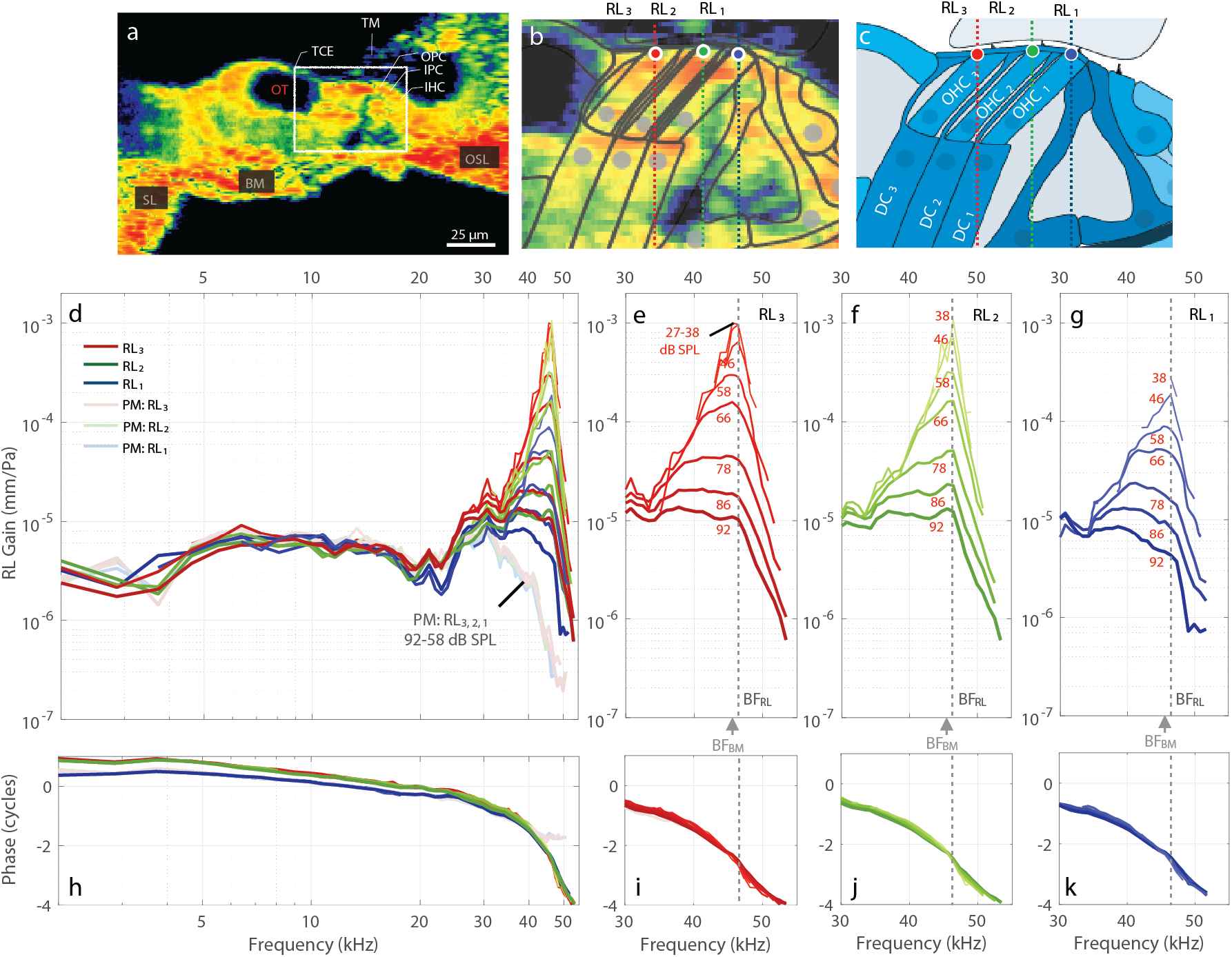
The in vivo and PM RL Gains across the apical ends of the three rows of OHCs (RL_3_, RL_2_, RL_1_; G637). **a** A 2D cross-sectional OCT image of a representative in vivo gerbil OoC with labeled key structures. Note that the OoC structure is displayed upside down in comparison to Fig. 1. **b** An enlargement of the region in panel **a** within the white box, with overlaid line drawings of the cells and other structures, and details of the measurement locations across the apical ends of the three rows of OHCs shown (RL_3_: red circle; RL_2_: green circle; and RL_1_:blue circle). Each colored vertical line indicates the lateral position and direction of the different OCT A-scans made during the OCT vibrometry measurements. **c** A labeled cross-sectional drawing from the 2D OCT image in panel **b**. **d** In vivo (dark red/green/blue colors) and PM (faded red/green/blue colors) gains for RL_3_, RL_2_, and RL_1_, respectively. **e–g** The in vivo near-BF gain region for RL_3_, RL_2_, and RL_1_, respectively, showing active amplification. The gains were highest at low SPLs (note that curves overlap for the 27–38 dB SPL stimuli in panel **e**. The overlaid numbers in panels **e–g** indicate the stimulus level for each gain response. **h–k** The phase responses corresponding to panels **d–g**, respectively. Note that in this figure the displacements are normalized by the sound pressure to produce gains in units of mm/Pa, and the phase responses are in units of cycles. The frequency axis is on a log scale in panels **d** and **h**, but on a linear scale in panels **e–g** and **i–k**.

**Fig. 4.**
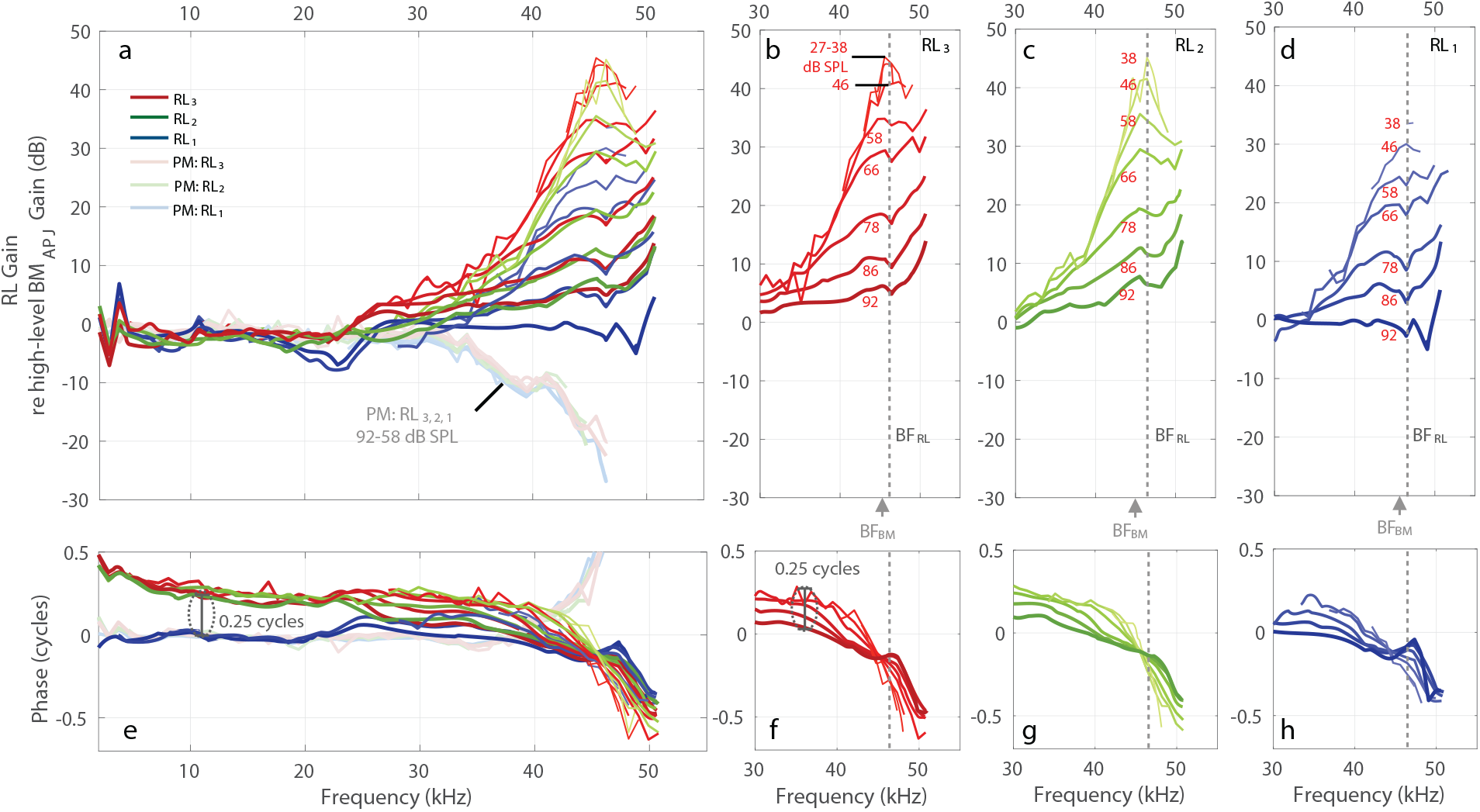
The in vivo and PM RL gains, normalized by the BM_APJ_ gain (measured at a high stimulus level), in order to remove the effects of the cochlear traveling wave, for specimen G637. **a** Direct comparisons of the in vivo and PM RL_3, 2, 1_ normalized gains (in dB). **b–d** In vivo normalized gains in the near-BF region for RL_3_, RL_2_, and RL_1_, respectively. **e–h** The phase responses corresponding to panels **a–d**, respectively. Note that the baseline high-level BM_APJ_ gain used for normalization was calculated as the average of ten measurements made at 92 dB SPL. All frequency axes in this figure are on a linear scale.

### Motion across the tops of the three OHC rows (RL_1–3_)

The motions in one animal at the points on the RL surface corresponding to the apical surface of each of the three rows of OHCs (RL_1_, RL_2_, and RL_3_) are shown in Figures 3–4. The gains in the living and dead animal, along with OCT images of the anatomy from the same living animal, are shown in Figure 3. To help remove the contribution to RL motion of the BM traveling wave, we normalized the RL gain by the BM gain at the junction between the arcuate and pectinate zones (BM_APJ_), by averaging multiple measurements made at a high stimulus level where there was relatively little cochlear amplification (Fig. 4). Results from additional animals are shown in Supplementary Information.

The data from all three RL rows show similar patterns of compressive nonlinear growth near BF, linear growth at low frequencies, and phases consistent with drive from a traveling wave (Fig. 3). In this animal, at frequencies near BF the motion amplitudes at RL_2_ and RL_3_ were similar, but the motion at RL_1_ was ~15 dB less (Figs. 3e–g, 4b–d). In contrast, below 20 kHz there was very little difference in the magnitudes of the RL gains, and furthermore, the magnitudes were similar in the living and dead animal (Figs. 3d, 4a). However, in the phase at low frequencies, RL_1_ had the same phase for the living and dead animal, whereas RL_2_ and RL_3_ had a phase advance of more than 0.25 cycles in the living animal compared to the dead animal or to RL_1_ (Fig. 4e).

Removing the traveling-wave phase (by normalizing by high-level BM_APJ_ motion) and plotting frequency on a linear scale allows the phase differences (and thus group delays) of RL_1–3_ to be seen more easily (Fig. 4). Near BF and down to 30 kHz, all three RL rows have phase patterns in which lower levels show a greater phase *advance* than higher levels (Fig. 4f–h; lower levels are thinner and have lighter colors). Above BF, this pattern reverses so that the phase *delay* is greater at lower levels. For all three RL rows, the group delays (the negative slopes of the phase-versus-frequency functions in Fig. 4f–h) become smaller as the sound level increases (Fig. 4f–h), which is consistent with the RL tuning becoming wider at higher sound levels. Interestingly, at each RL row the phase-versus-frequency curves from different sound levels cross at a frequency that is slightly lower than BF (like the pattern reported for BM motion; Robles and Ruggero, 2001). The phase-crossing frequency is similar in RL_2_ and RL_3_, but slightly lower in frequency for RL_1_ (Fig. 4f–h). Another interesting feature of the data in Figure 4 is that at high levels the RL gain relative to the high-level BM_APJ_ gain peaks at a frequency that is above the BFs of the BM and RL (arrowheads versus dashed lines in Fig. 4). We will return to this in the Discussion.

The RL gain relative to the high-level BM_APJ_ gain, compared across animals, is shown in Figure 5. Each horizontal-axis position shows summary statistics as box plots and individual data points as symbols from each animal, in the same animal order in all panels (and in later summary figures). From left to right, each column summarizes data from a different frequency, with the magnitudes shown in the top row and phases shown in the bottom row. The results in all columns were measured using the highest stimulus level common to measurements for all of the structures in any given animal, except for those in the third column (BF), which were measured using the lowest level available. At a representative low frequency (10 kHz), all but one animal has RL gains within 10 dB of the high-level BM_APJ_ gain, and most RL_3_ gains are higher than the RL_1_ gains (Fig. 5a). All of the RL_2_ and RL_3_ phases are advanced relative to BM_APJ_ by close to 0.25 cycles, but 7 of 9 RL_1_ phases are close to the BM_APJ_ phase (Fig. 5e). At 30 kHz (Fig. 5b, f), which is close to 0.5 octaves below the BFs, most RL gains are within 5 dB of the high-level BM_APJ_ gain, and the phases are in a narrow range between 0 and 0.25 cycles of the BM_APJ_ phase, with all of the RL_1_ phases less than the RL_3_ phases (Fig. 5b, f). At BF (Fig. 5c, g) there are a wide range of gains across animals, but in all cases the gain of RL_3_ is significantly greater than that of RL_1_, by 10 ± 1 dB (p <0.001, Fig. 5c). At BF, the RL_1–3_ phases are mostly within 0.25 cycles of the BM_APJ_ phase, and in each animal (except one), the RL_1_ and RL_3_ phases are close. At the above-BF frequency, measurements of RL motion relative to BM_APJ_ motion (which were usually done at the highest frequency for which reliable measurements were available in the high-level BM_APJ_ gain used for normalization), the RL gains are consistently greater than the BM_APJ_ gain and in almost all cases exhibit a phase lag compared to the BM_APJ_ phase (Fig. 5d, h).

**Fig. 5.**
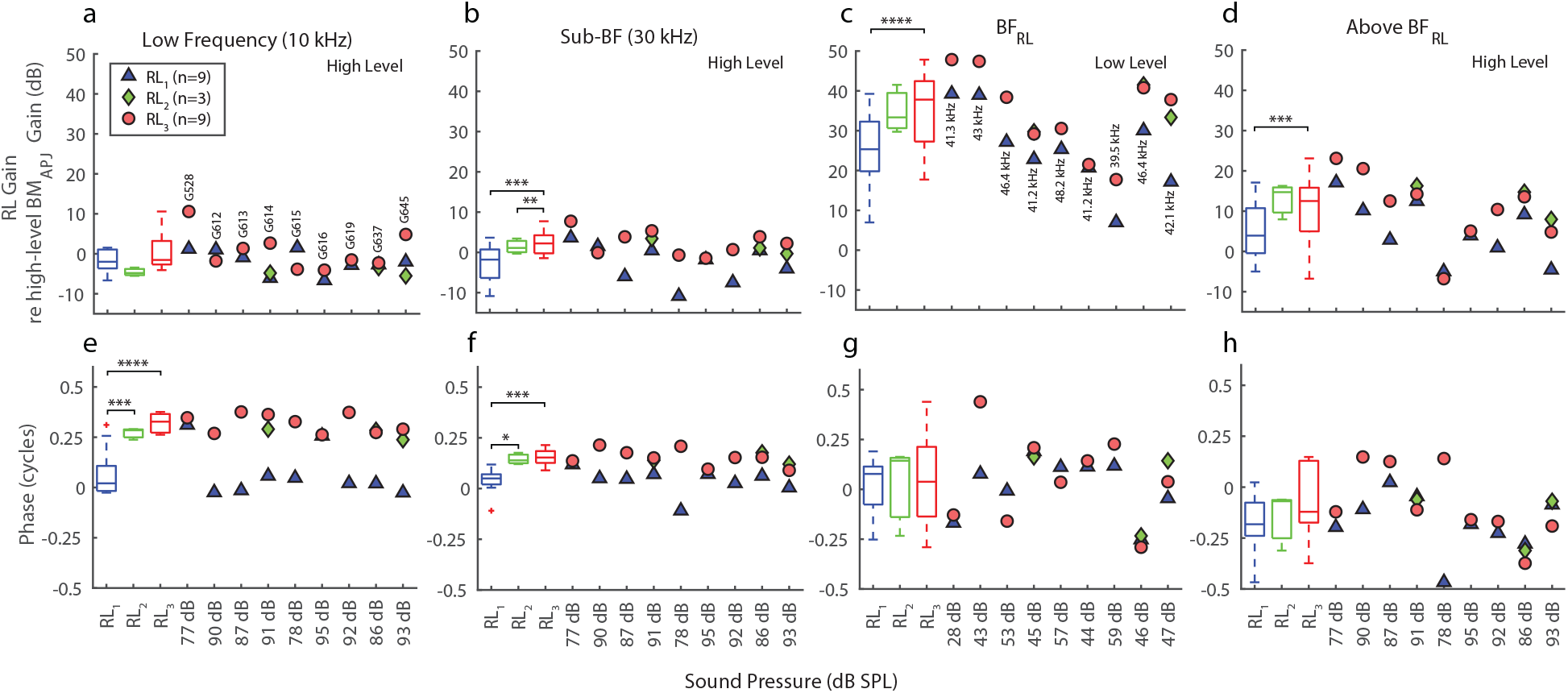
Summary comparisons across animals of the in vivo normalized gains at the apical ends of the OHC rows (RL_3, 2, 1_), for selected frequencies. **a–d** The respective normalized-gain magnitudes at a low frequency (10 kHz, high stimulus level), near 30 kHz (Sub-BF, high stimulus level), at BF_RL_ (low stimulus level), and above BF (high stimulus level). The above-BF_RL_ frequencies used in panels **d** and **h** correspond to the high-stimulus-level magnitude peak for each animal. **e–h** The phase summaries corresponding to panels **a–d**, respectively. The individual datapoints in columns 1–2 were averaged over a 1/3-octave width centered around 10 and 30 kHz, respectively, while those in column 4 were averaged over a 1/6-octave width centered around the selected frequency. The points shown in column 3 each represent the value at a single frequency. Note that the stimulus levels listed on the horizontal axes and BF values labeled in panel **c** vary across animals (labeled in panel **a**). The box-and-whisker plots on the left-hand side of each panel provide statistical summaries of the individual results, matched by color. The ANOVA *p*-values between categories are indicated by horizontal brackets at the top. The number of asterisks above each bracket corresponds to the following degrees of statistical significance: *p* < 0.05 (*), *p* < 0.02 (**), *p* < 0.006 (***), and *p* < 0.001 (****), all of which are of greater significance than the *p_0_* = 0.05 criterion. Insignificant *p*-values are not displayed. The red ‘+’ symbols indicate outliers, the boxes indicate the interquartile range (IQR) with 95% confidence, and the horizontal line inside each box indicates the median value. The whiskers indicate the minimum and maximum of the range (excluding outliers). In some cases the RL_2_ whiskers are too small to be visible. The RL_3, 2, 1_ results for animal G637 are shown in Figs. 3 and 4.

### Motion of the three OHC rows at their junctions with the Deiters’ cells (OHC-DC-junction-_1–3_)

The motions in one animal across the bottoms of the three rows of OHCs at their junctions with the Deiters’ cells (OHC-DC-junction_1–3_) are shown in Figure 6. The OHC-DC-junction gains (the displacement divided by the sound pressure) are shown on the left with a log-scaled frequency axis, and the OHC-DC-junction gains with the traveling-wave drive removed (through normalization by the high-level BM_APJ_ gain) are shown on the right with a linear frequency axis. The gains and phases at selected frequencies are shown for all animals in Figure 7. The OHC-DC-junction data vary relatively little across animals, and the observations for the example animal (G637) generally hold across all animals (Fig. 7).

**Fig. 6.**
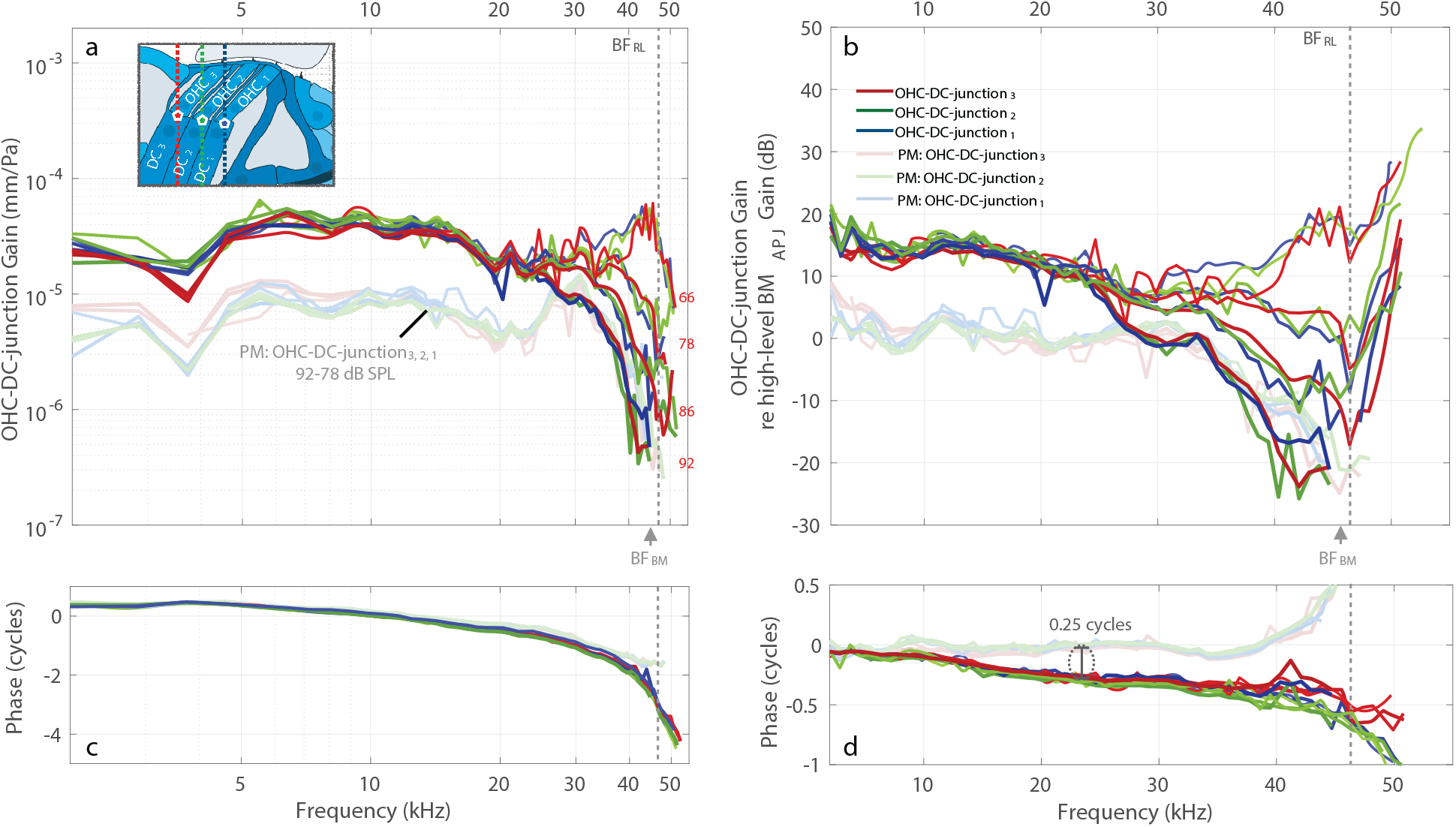
Motion comparisons of the OHC-DC junctions at the basal ends of the OHCs, across the three OHC rows (OHC-DC-junction_3, 2, 1_), for specimen G637. **a** In vivo (darker colors) and PM (faded colors) OHC-DC-junction_3, 2, 1_ gains relative to the sound pressure (in units of mm/Pa). **b** The gains in **a** normalized by the high-level (92 dB SPL) BM_APJ_ gain (in dB). Note that the inset drawing in panel **a** shows the OCT-vibrometry measurement locations for OHC-DC-junction_3_ (red pentagon), OHC-DC-junction_2_ (green pentagon), and OHC-DC-junction_1_ (blue pentagon). Each colored vertical line indicates the lateral position and direction of the different OCT A-scans acquired during the OCT vibrometry measurements. **c–d** The phase responses corresponding to panels **a–b**, respectively. Note that the frequency axes are on a log scale in the first column and on a linear scale in the second column.

**Fig. 7.**
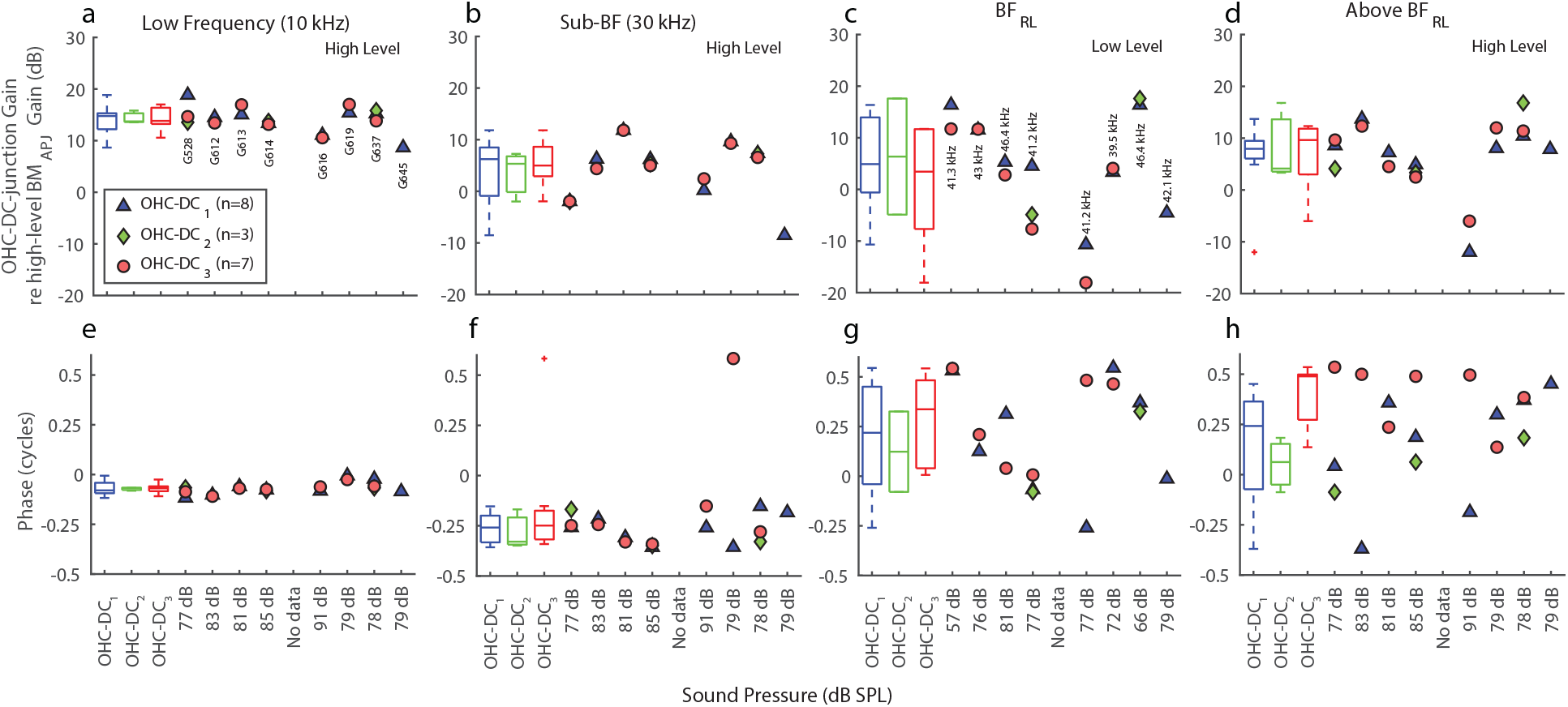
Summary comparisons of the in vivo normalized OHC-DC-junction at the basal ends of the OHCs gains across animals for the three OHC rows (OHC-DC_3, 2, 1_). **a–h** The normalized-gain magnitudes (**a–d**) and corresponding phases (**e–h**), each with respective box-and-whisker-plot summaries, using the same frequencies and methods described in Fig. 5. The ANOVA *p*-values are all insignificant in this figure. The OHC-DC-junction_3, 2, 1_ results for animal G637 are shown in Fig. 6.

In the example animal, there is little difference in the gains of the three OHC-DC-junction rows at any frequency or level (Fig. 6a, b). At frequencies near BF, all three OHC-DC-junction rows have compressive growth and are similar in gain (Fig. 6). Near BF, the three OHC-DC-junction rows have broad peaks at low sound levels, but at higher levels the amplitude decreases as the frequency increases, and above BF it tends to decrease (Fig. 6a). At frequencies near BF and at low sound levels, the OHC-DC-junction gain is greater than the high-level BM_APJ_ gain, but at high sound levels the OHC-DC-junction gain is less than the high-level BM_APJ_ gain (values below zero dB in Fig. 6b). At frequencies just below BF (e.g., 40 kHz), the gain of the OHC-DC-junction motion at the highest stimulus level is actually less than in the dead animal (Fig. 6b), which implies that the active motion was partially cancelling the passive motion. A similar pattern is seen in the other animals (Supplementary Information Figures 6B, 7B).

At low frequencies (an octave or more below BF), the three OHC-DC-junction rows show linear growth at the levels tested, and considerable gain (~15 dB) relative to the OHC-DC-junction gain in the dead animal (light color lines in Fig. 6a) and relative to the high-level BM_APJ_ gain (Fig. 6b). Above BF, as the frequency increases, the OHC-DC-junction gain decreases relatively little (Fig. 6a), but the BM_APJ_ gain decreases faster so that above BF the OHC-DC-junction gain is greater than the high-level BM_APJ_ gain (above BF the lines go up in Fig. 6b).

The phases of the OHC-DC-junction gains are similar across the three rows at mid frequencies, but the phases deviate from one other at high and low frequencies (Fig. 6c, d). At mid frequencies the OHC-DC-junction phases lag behind the high-level BM_APJ_ phase and the OHC-DC-junction phase of the dead animal by ~0.25 cycles (Fig. 6d). At the lowest frequencies, the OHC-DC-junction phase lags approach zero (Fig. 6d). Near the BM_APJ_ BF, the OHC-DC-junction_3_ phase lags behind the high-level BM_APJ_ phase by ~0.25 cycles, with rows 1 and 2 lagging more.

### Motion along the basilar membrane

The gains and phases in the example animal at three points along the width of the BM, i.e., in the arcuate zone (BM_AZ_), at BM_APJ_, and in the pectinate zone (BM_PZ_), are shown in Figure 8. The locations of BM_AZ_, BM_APJ_, and BM_PZ_ are shown in the inset of Figure 8a. The gains and phases at selected frequencies are shown for all animals in Figure 9.

**Fig. 8.**
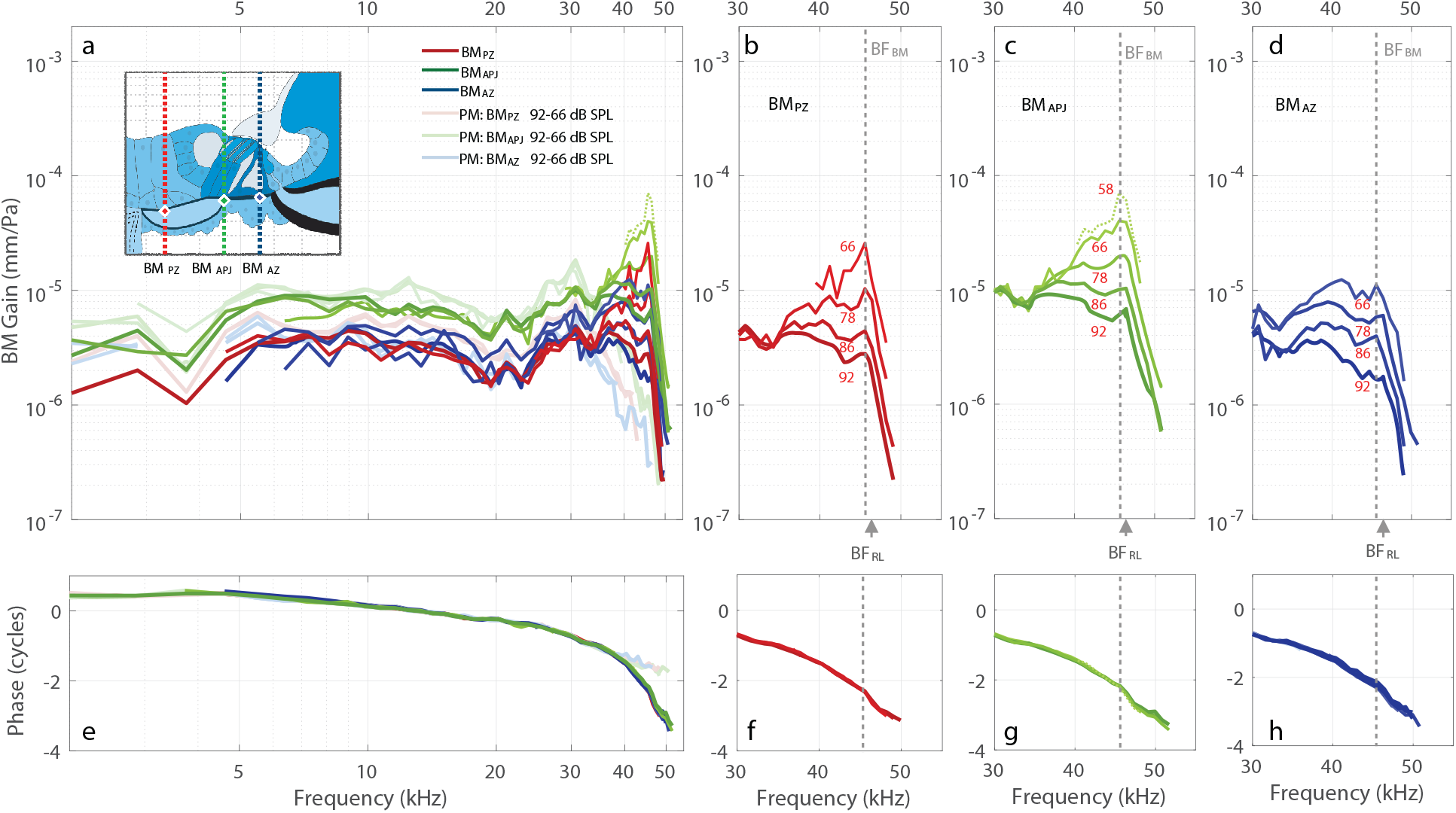
In vivo and PM motions across three BM locations (BM_PZ_, BM_APJ_, and BM_AZ_), for specimen G637. **a** In vivo (darker colors) and PM (faded colors) BM_PZ_, BM_APJ_, and BM_AZ_ gains with respect to the stimulus pressure (in units of mm/Pa). The green dotted line indicates the lowest-level BM_APJ_ measurement. The inset drawing shows the measurement locations for BM_PZ_ (red diamond) BM_APJ_ (green diamond), and BM_AZ_ (blue diamond). Each colored vertical line indicates the lateral position and direction of the different OCT A-scans acquired during the OCT vibrometry measurements. **b–d** The respective in vivo gains of the different BM locations, showing active nonlinear amplification in the near-BF region. **e–h** The respective phase responses corresponding to panels **a–d**. Note that the displacements are normalized by the sound pressure in all panels. The frequency axes in the first column are on a log scale, and those of the remaining columns are on a linear scale.

**Fig. 9.**
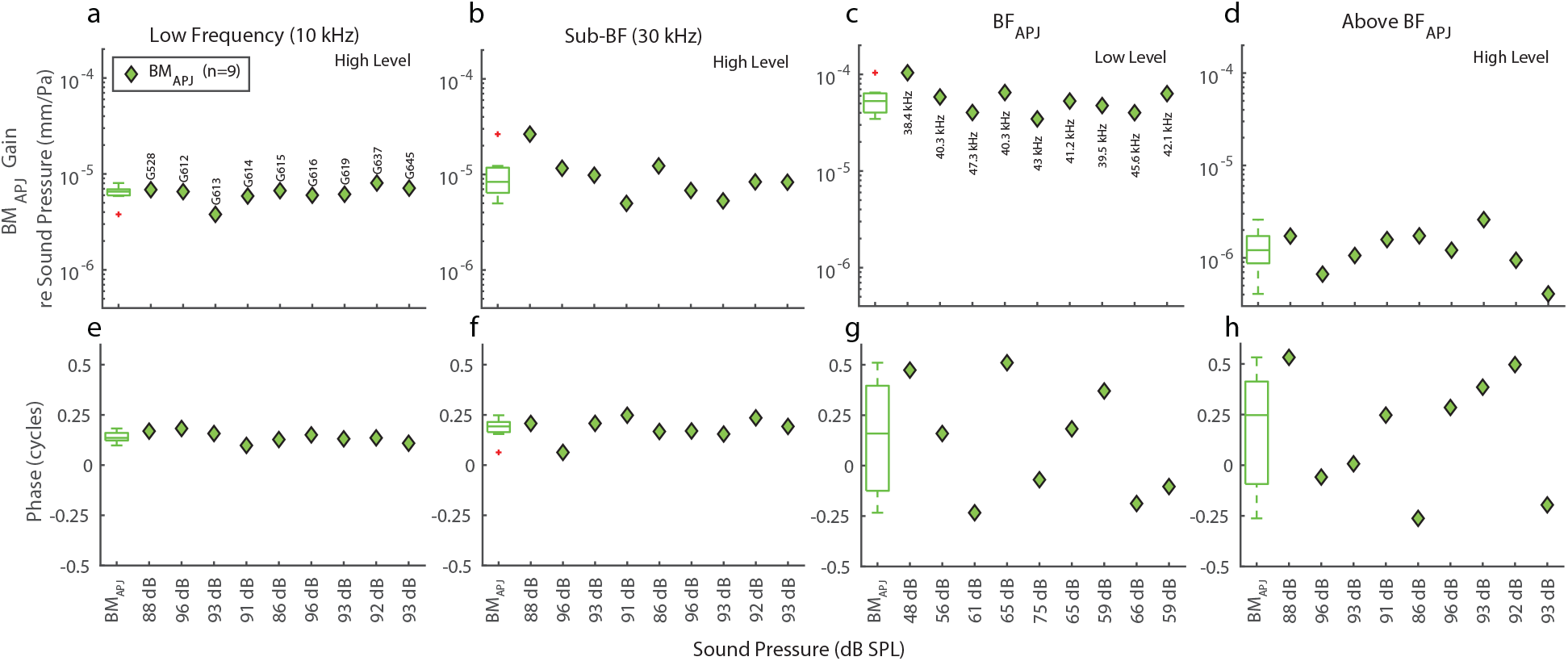
Summary comparisons across animals of the in vivo BM_APJ_ gains with respect to the sound pressure (in units of mm/Pa), for selected frequencies. **a–h** The gain magnitudes (**a–d**) and corresponding phases (**e–h**), each with a respective box-and-whisker-plot summary. The frequencies in the first two columns match those of Figs. 5 and 7, hover those used in column 3 correspond to BF_BM_ (as shown in Fig. 8) instead of BF_RL_, and those used in column 4 are quarter-octave above the frequencies used in column 3. The BM_APJ_ results for animal G637 are shown in Fig. 8.

As expected from previous work, the largest BM gain is at the arcuate-pectinate junction, and the phase versus frequency of the BM gain exhibits a phase delay that increases as frequency increases, consistent with excitation by a traveling wave. Near BF the BM gain grows compressively, and below BF the growth of the BM gain is mostly linear.

## Discussion

Our measurements present a different and more complex picture of the motion within the OoC than has been conveyed by previously published measurements in the gerbil. We were able to make more-detailed measurements because our OCT system has better spatial resolution than the systems used in previous papers (Ren et al., 2016; Ren and He, 2020; He et al., 2018; He and Ren, 2021; Fallah et al., 2019; 2021; Strimbu et al., 2020; Strimbu and Olson, 2021; Cooper et al., 2018). In previous publications, the OCT images had comparatively lower resolutions, and although one could identify the BM as the first region of reflectivity past the scale, other structures within the OoC could not be identified based on their shapes in the image. Instead, the locations of the inner and outer tunnels were identified and the locations of structures such as the OHCs and/or the RL were inferred from the locations of the tunnels (Ren and co-workers did not have images and just used distances from the BM). In contrast, our images are of sufficiently high quality to allow us to identify the RL and the bottoms (base) of the OHCs (Fig. 1e) without needing to determine their positions based on an overlaid standard image or estimates of where reflectivity measurements originated from. In some cases, three reflective peaks could be seen on the RL at the tops (apical ends) of the three rows of OHCs. We identified measurements at different radial locations according to the OHC row they were closest to, but the alignment may not have been exact. For instance, we do not know if the structures that produce the reflectivity peaks on the RL correspond to an OHC top or an adjacent PhP top. Despite this uncertainty, our ability to discern radial positioning along the OHC rows is far greater than any previous publication, and this is the first publication to definitively identify the RL in the gerbil basal region and to measure from it.

### Our data compared to previous reports

Previous reports used a variety of optical-beam wavelengths, bandwidths, and motion-detection processing, and since the reflectivity of a structure depends on the wavelength, motion from different structures is likely to have been measured in different experiments. In addition, the motion measured depends on the viewing angle, in part because different viewing angles can change the reflectivity of a structure, possibly due to birefringent properties of collagen fibrils in cochlear structures (Kalwani et al., 2013). Further, if the OoC is not viewed perpendicular to the BM, then supposedly “transverse” measurements will also include radial and longitudinal components. For instance, Cooper et al. (2018) measured from the gerbil 20-kHz region as viewed through the round-window membrane (RWM). To view the 20-kHz region through the RWM, the OCT beam must be pointed tangentially toward the apex (Fig. 1c), such that the motion measured along the axial direction of the beam will be a combination of transverse and longitudinal motion. By changing the angle of their beam, Cooper et al. (2018) concluded that much of what they measured was longitudinal motion. We developed a gentle surgical technique that produces little trauma and allowed us to measure in the 40–50-kHz BF region. Measurements in this region allowed us to use an OCT beam angle that is almost perpendicular to the BM when viewed through the RWM (Fig. 1c). Thus, our measurements are dominated by the transverse vector component of motion, with little contribution from radial or longitudinal components.

Motion measurements from within the OoC in the gerbil base have been reported by three other groups (Ren et al., 2016; Ren and He, 2020; He et al., 2018; He and Ren, 2021; Fallah et al., 2019; 2021; Strimbu et al., 2020; Strimbu and Olson, 2021; Cooper et al., 2018). Our measurements differ from previous reports in that: (1) we measured at three transverse locations, i.e., the BM, the OHC-DC-junction and the RL, whereas previous measurements were only at two locations, the BM and in the “OHC region”; and (2) we measured at several locations radially across the OoC at points aligned with the three rows of OHCs. There are no previous systematic measurements radially across the OHCs. We first consider the measurements at different distances from the BM; later we consider measurements at different radial locations. We, and all previous reports, found the classic pattern of BM motion: in the BF region the BM motion was amplified (relative to the dead animal) and had compressive growth. At frequencies 0.5 octaves or more below the BF, the BM response was not amplified and grew linearly with level.

In the gerbil base, we found motions within the OoC that are substantially different from the motions previously reported (Ren et al., 2016; Ren and He, 2020; He et al., 2018; He and Ren, 2021; Fallah et al., 2019; 2021; Strimbu et al., 2020; Strimbu and Olson, 2021; Cooper et al., 2018). The primary reason for this difference appears to be that we measured from different structures than previous reports and/or used different viewing angles. We measured motion at the RL and OHC-DC-junction and identified those regions from images that were clear enough that we can be confident that those are the structures that were measured (Fig. 1, Fig. 3a). The images from the other reports did not allow such definitive identification of the structures producing the measured reflections, and the measured structures were referred to loosely as the “OHC region” (Olson group’s measurements), the “hot-spot” in the OHC-Deiters’ cell region (Cooper et al., 2018) or as the “RL” (Ren group’s measurements, although “RL” was a blind identification that the Olson group in Fallah et al., 2019, interpreted as corresponding to their “OHC region”). We will refer to the non-BM measurements from the previous reports as “OHC region” measurements. A simplification that may be fairly accurate is that the “OHC region” motions reported by others show a combination of the motions we measured at the OHC-DC-junction and RL points.

For frequencies near BF, the previously reported “OHC region” motions show compressive nonlinearity and more gain than the BM motion. We found compressive nonlinearity near BF in both the OHC-DC-junction and RL motions. Near BF, our RL motion looks like the reported “OHC region” motion and usually has more gain than the BM motion (for RL_2_ and RL_3_, but not always for RL_1_ at the highest level; Fig. 4). In contrast, the OHC-DC-junction motion near BF is generally less than the BM motion (compare Figs. 6a and 9c), and has a substantially different pattern versus frequency than the reported “OHC region” motions.

For frequencies more than 0.5 octaves below BF (low frequencies), the previous reports found the “OHC region” motions to be substantially more (10–20 dB) than the BM motion in the live animal, but similar to the BM motion in the dead animal, which shows that the “OHC region” motions received substantial amplification at low frequencies. Our low-frequency OHC-DC-junction motion is also 10–20 dB more than the BM motion in the live animal, and similar to the BM motion in the dead animal. Thus, at low frequencies our OHC-DC-junction motions are similar to previously reported “OHC region” motions. In contrast, at low frequencies our BM_APJ_-normalized RL gain is seldom very different from 0 dB (Figs. 4, 5a). Interestingly, the low-frequency motion of RL_1_ has a phase similar to the BM_APJ_ motion (Fig. 5e), but RL_2_ and RL_3_ have phases that are advanced by about 0.25 cycles at the lowest frequencies (Fig. 5e). Our data show that at low frequencies the RL gain magnitude is similar to the BM_APJ_ gain magnitude (Fig. 5a), but the OHC-DC-junction motion (which is between the RL and the BM) is 10–20 dB greater (Fig. 7a)!

### RL motion *above* BF

One interesting finding is that at frequencies above BF, the RL gain is larger than the BM gain (Figs. 4, 5; Supplementary Information Figs. 1–3). This is due to the BM having a slightly lower BF than the RL, and to the BM response falling very sharply as frequency increases, starting at frequencies just above the BF_BM_, whereas the RL motion does not fall as sharply (Fig. 10, a versus c). Similar findings were made by Ren et al. (2016) in gerbils and Chen et al. (2011) in guinea pigs. One explanation for this is that traveling-wave cochlear amplification falls rapidly above BF, which decreases BM motion rapidly. RL motion rides on top of BM motion and also decreases, but RL motion above BF also receives local amplification by the OHCs, just as it does below BF. Since the OHC stereocilia are no longer saturated by the drive from BM motion, they can amplify RL motion more than at BF. This is consistent with the effects seen by applying salicylate or furosemide (Strimbu et al., 2018; 2020).

**Fig. 10.**
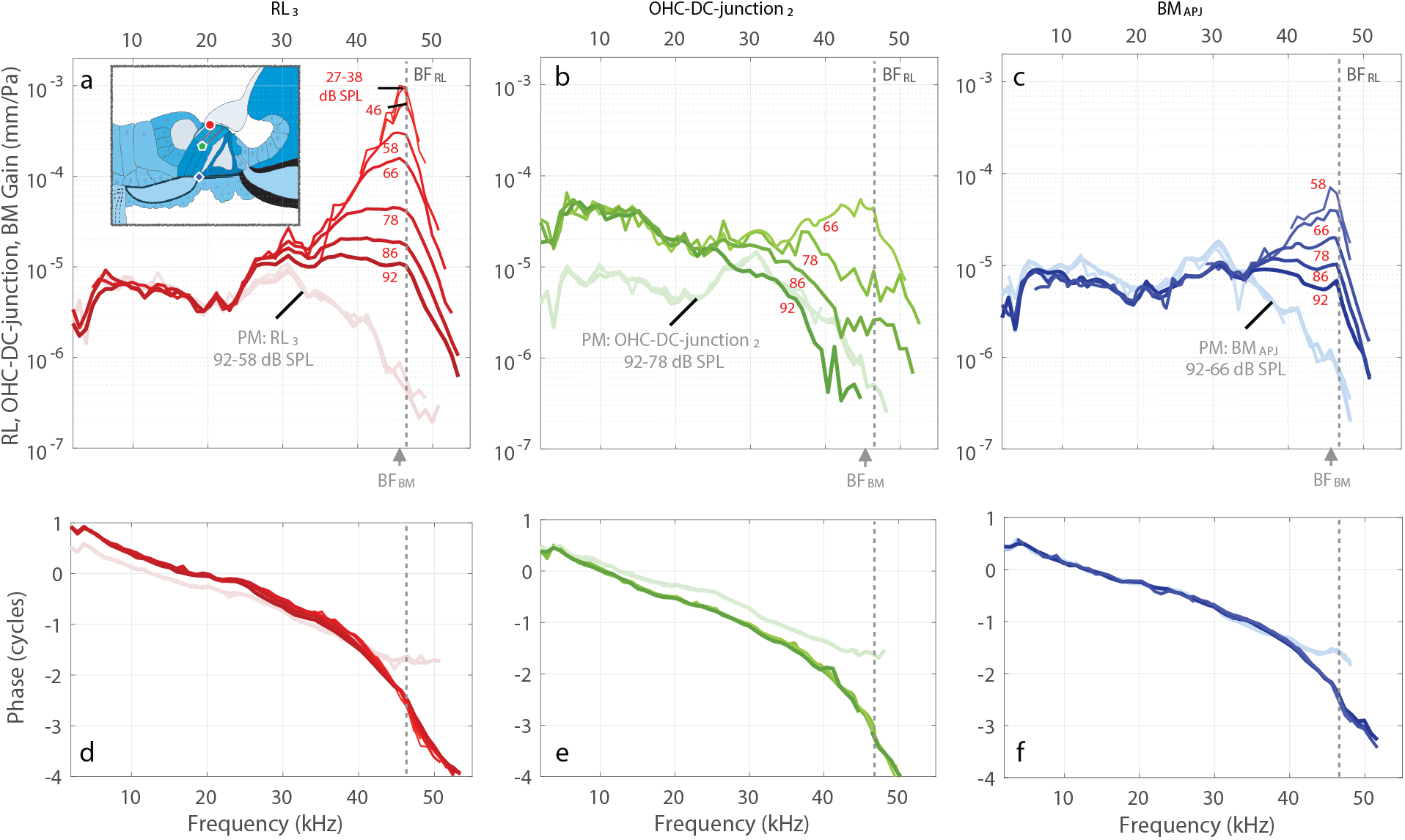
In vivo (darker colors) and PM (faded colors) gain comparisons across different OoC structures along a transverse direction, for specimen G637. **a–c** The respective gain magnitudes of RL_3_ (red), OHC-DC-junction_2_ (green), and BM_APJ_ (blue), all relative to the sound pressure (in units of mm/Pa). The inset drawing in panel **a** shows the measurement locations for RL_3_ (red circle), OHC-DC-junction_2_ (green pentagon), and BM_APJ_ (blue diamond). **d–f** The respective phase responses corresponding to **a–c**. Note that the available stimulus levels vary across the structures due to different signal-to-noise ratios.

### Motion at different radial positions within the organ of Corti

This is the first report of motion within the OoC across the three OHC rows, either at the top or the bottom of the OHCs. For frequencies near BF, RL_3_ has significantly more motion (10 ± 1 dB) than RL_1_ (Fig. 5c), but at the OHC-DC-junction there is relatively little difference across the three OHC rows (Fig. 7c). For frequencies below BF, the RL-gain magnitudes are similar across the three rows but the phase of RL_1_ is about 0.25 cycles lower than that of RL_2_ and RL_3_ (p<0.006) (Fig. 5a), while for OHC-DC-junction the gain magnitudes and phases are similar across the three rows (Fig. 7a). For the RL_1_ and RL_3_ motions to differ significantly from one another while the OHC-DC_1_ and OHC-DC_3_ motions do not differ seems to imply that the RL mosaic comprised of OHC and PhP cuticular plates and adhesion molecules is much more flexible than the DCs connected to the OHCs at their bottom surfaces.

Two previous reports provide sparse measurements from live animals that are consistent with our seeing more motion at RL_3_ than RL_1_. Fallah et al. (2019) reported measurements done at two radially separated “OHC regions” and found more motion in the more-lateral location in one case, but not in the other. Dewey et al., (2021) reported one set of interpolated measurements along the top of the mouse apical region OoC and found similar motions at RL_1_ and RL_3_ with 5 dB greater motion in the middle near RL_2_. Also relevant are measurements in an excised gerbil middle-turn preparation that found differences in motion in the radial and longitudinal directions across the three OHC rows at the bottom of the OHCs with electrical stimulation (Karavitaki and Mountain, 2007).

What do the differences in the motions across RL_1_, RL_2_, and RL_3_ tell us about what is happening in the cochlea? Across our animals with available measurements, the motions of RL_2_ and RL_3_ are similar and those of RL_1_ are different (Figs. 3–5). At high frequencies RL_2_ and RL_3_ have higher gains than RL_1_, but all have similar phases, while at low frequencies they all have similar gains but RL_2_ and RL_3_ have ~0.25-cycle phase advances relative to RL_1_. There are two (not mutually exclusive) classes of interpretations for the motion differences: (1) the RL motions are transverse motions with little contribution from radial and/or longitudinal RL motion, and (2) the differences are primarily from non-transverse motions (radial and/or longitudinal). One simple way for there to be more transverse motion at RL_3_ than RL_1_ is for these motions to be made up of an OoC transverse motion plus a transverse rotation of the RL about a fulcrum near RL_1_ (e.g., at the top of the pillars as suggested by Nowotny and Gummer, 2008; 2011). Our RL data might be matched if the rotation and translation were in-phase near BF, and if the rotation had a phase advance at low frequencies. Counting against the RL-rotation hypothesis is its implication that the RL_2_ motion should be about half-way between the RL_1_ and RL_3_ motions, while the RL_2_ motion is instead similar to the RL_3_ motion. However, there are not enough RL_2_ measurements to rule out the rotation hypothesis. If RL rotation is not a big factor, and the motion is all transverse, this would mean that there is significant deformation of the RL.

Another possibility is that the RL_3_ and RL_2_ motions are greater than those of RL_1_ because of non-transverse motion. This might explain the differences observed at low frequencies, because a phase difference of ~0.25 cycles can come from combining motions that are occurring in two perpendicular directions (Cooper et al., 2018). Also, the motions at RL_1_ and BM_APJ_ can be expected to be similar because these two regions are connected by the relatively stiff outer pillar cells. However, near BF the RL_3_/RL_1_ ratio is about 10 dB or a factor of 3. To account for this large difference in motion there would have to be almost an order-of-magnitude more non-transverse motion than transverse motion at RL_3_ compared to RL_2_, because viewing angle is nearly transverse and would record only a small fraction of any radial or longitudinal motion. An order of magnitude more non-transverse motion than transverse motion in RL_3_, but not in RL_1_, seems unlikely. This does not mean that the RL_2_ and RL_3_ measurements do not include any non-transverse contributions, but that non-transverse contributions are unlikely to account for the observed differences between RL_3_ and RL_1_. One conclusion from all of this is that to account for our measurements, it is likely that there is substantially more transverse motion of RL_3_ compared to RL_1_, which means that there must be substantial deformation of the RL. In the mouse, Dewey et al. (2021) reported finding a reversal of the motion along the top of the OoC in the region next to RL_3_ at the attachment of the PhP furthest from the pillar cells, which would be consistent with this part of the RL being not very stiff.

One implication of the interpretation that the increased motions of RL_2_ and RL_3_ compared to RL_1_ are due to transverse motion, is that the OHCs produced more motion at RL_2_ and RL_3_ than at RL_1_. This could be because all OHC rows receive the same stereocilia deflections and produce the same OHC force, but that RL_1_ motion is restrained by its close attachment to the pillar cells. Alternately, the OHCs of rows 2 and 3 might receive larger stereocilia deflections and therefore produce more force. This would be consistent with there being more radial motion at RL_2_ and RL_3_, although this radial motion might contribute little to the transverse motion that we measured. Recently it was hypothesized that this increased radial motion could originate from expansion or contraction of the outer tunnel, whose upper wall connects to the RL near RL_3_ via the tectal-cell extension (Cho et al., 2022). Measurements from two or more angles are needed to decompose the motion directions into vector components at RL_1–3_ to test these possibilities. Another approach would be to use a finite-element model of the RL mosaic that incorporates the radial and longitudinal angles of the PhPs across the three OHC rows, as well as the outer-tunnel fluid space. Regardless of the underlying reason(s) for the observed motion differences, these measurements suggest that the RL may not move as a stiff plate hinging around the pillar-cell heads as has been assumed, but that its mosaic structure may instead bend and/or stretch. Understanding the specifics of RL motion is fundamental to understanding both cochlear amplification via OHC stimulation and sound transduction via inner-hair-cell stimulation.

### Do we see a motion correlate for the frequency at which cochlear amplification starts?

The phasing of OHC motion to amplify BM motion arises from a change in the phase of RL motion that, as frequency is increased, begins rather abruptly at about 0.5 octaves below BF (Dong and Olson, 2013; Lee et al., 2016; Guinan, 2020). Since the Dong and Olson (2013) measurements that identified this rapid phase change were done in the gerbil base, it seemed possible that a motion correlate of this transition would become evident in our measured motions. We have looked for such a correlate but have not been able to identify any change in the motion, such as a bend in the curves, that occurs around 0.5 octaves below BF. This could be because our RL measurements are almost purely transverse and are therefore insensitive to changes in RL radial motion.

## Methods

### Animal preparation

Healthy female Mongolian gerbils, (N=19, aged 5–11 weeks, weight range 41–76 g) were used. Gerbils have the advantage of being small and having been previously used in experiments on intra-cochlear pressure (Olson, 1999, 2001; Kale and Olson, 2015), and in OCT cochlear measurements (*e.g*., Dong et al., 2018; Cooper et al., 2018; Fallah et al., 2019; 2021; Strimbu et al., 2020; Strimbu and Olson, 2021).

To minimize hearing damage, care was taken throughout to reduce noise and vibration (Brown, 1983). A key surgical technique was opening the bulla after applying phosphoric acid gel (PAG, PULPDENT Corporation, MA, USA) to thin (decalcify) and soften the bone (Alyono et al., 2015). PAG was applied for 10 seconds and then wiped off using a micro-wipe tip. After repeating this three times, a narrow opening in the posterolateral wall of the bulla was made with fine forceps. The tympanic membrane, malleus, incus, stapes, and RWM were all kept intact.

Surgical procedures and anesthesia protocols were described previously (Cho et al., 2022). After the in vivo measurements, an intraperitoneal injection of Fatal Plus (>150 mg/kg) produced euthanasia. Within 5–10 minutes after the injection, the heart typically stopped and the animal stopped breathing. The postmortem vibration measurements were done 5 to 60 min after the animal stopped breathing and had no heartbeat. This study was approved by the Institutional Animal Care and Use Committee (IACUC) at Massachusetts Eye and Ear.

### Stimulus generation and acoustic system

Signal generation and sound-pressure control used a National Instruments (NI) PXI-4461 dynamic signal acquisition board mounted in an NI PXI-1031 chassis with an NI PXI-8196 embedded computer (NI, TX, USA). For measuring distortion product otoacoustic emissions (DPOAEs) this was controlled using the LabVIEW-based Cochlear Function Test Suite (EPL_CFTS, version 2.37-R3041; Hickman *et al*., 2021). For the OCT vibrometry measurements, this used custom stimulus generation and synchronous measurement software (SyncAv, version 0.42; Gottlieb *et al*., 2016), which generated a sequence of pure tones (2–63 kHz, ~0.8 kHz linear frequency steps). The board output was amplified by a Techron Model 5507 power amplifier (AE Techron, IN, USA) which drove a Parts Express 275-010, tweeter speaker mounted in a custom-built closed-field acoustic assembly. During the experiments, ear-canal pressure was measured by a calibrated Knowles FG-23329 electret microphone and probe tube with the probe-tube opening placed 1–2 mm from the tympanic membrane. Initially, an in situ pressure response with a constant stimulus voltage as a function of frequency was measured. The measured microphone response was then used to vary the stimulus voltage to equalize the ear-canal pressure and produce a near flat response across frequencies (Fig. 2a). This equalization voltage curve was scaled in approximately 10 dB steps (or ~5 dB steps at higher levels) to produce the range of stimulus levels measured in the experiment. The value for the stimulus sound pressure level (SPL; in dB) associated with each measurement was calculated as the average measured pressure across frequencies (0 dB SPL = 20 μPa). The stimuli were varied from ~30 to ~90 dB SPL across experiments, and the actual SPLs varied ±4 dB due to small differences in the position of the acoustic assembly in the ear canal from the initial equalization steps.

### Monitoring cochlear sensitivity

Cochlear sensitivity was monitored by DPOAEs done before and after surgery, and approximately every 20 min during in vivo measurements using the EPL_CFTS software (Hickman *et al*., 2021). DPOAEs were evoked by a series of tones at frequencies f1 and f2 (f2/f1 = 1.2), with f2 varied from 2 to 63 kHz in 0.5-octave steps below 30 kHz, and 2-kHz steps above 30 kHz. The tones were presented at equal levels in separate runs at 50 or 70 dB SPL. Across animals, the DPOAE amplitudes mostly ranged from ~15 to 20 dB SPL for f2 frequencies above 30 kHz. A cochlear region’s sensitivity was assessed from its DPOAE with f2 near BF, and was deemed adequately sensitive if the DPOAE from 50 dB SPL primary tones was greater than 10 dB SPL and the noise floor was less than 3 dB SPL.

### OCT Imaging and Vibrometry

All OCT imaging and vibrometry measurements were done using an SD-OCT system with a 900-nm center wavelength and a high-speed (up to 240-kHz) line-scan camera (GAN620C1, Thorlabs, Germany). The OCT system combined two nearinfrared superluminescent-diode light sources with center wavelengths of 847.5 nm and 929.6 nm, with full-width, half-maximum (FWHM) spectral bandwidths of 63.4 and 95.8 nm, respectively. Before performing the fast Fourier transform (FFT), the SD-OCT system multiplied the recorded interference spectrum with an apodization (Hann) window function, which caused a broadening of the axial beam profile by a factor of 2 and had an impact on the axial resolution. The penetration depth was ~1.44 mm (in water, with a refractive index of 1.33). The axial resolution was ~2.23 μm (in water) and the lateral resolution was ~8 μm, using a 36-mm, 0.055-NA, 2× objective lens (OCT-LK3-BB, Thorlabs, Germany).

The OCT measurements were done with custom LabVIEW (NI, TX, USA)-based VibOCT software (version 2.1.4), built using the Thorlabs SpectralRadar software development kit (version 5.4.8). This system can provide (1) real-time video images that were used in the anatomical approach to determine the region of interest for OCT scans; (2) depth-resolved 1D A-scans (reflectivity vs. axial depth, e.g., Fig. 1d); (3) 2D cross-sectional B-scans (axial depth vs. scan range, e.g., Fig.1e) with software x-y offset and scan angle; (4) 3D volumetric C-scans (reflectivity vs. axial depth with scan range over two perpendicular axes), and (5) synchronous vibrometry data acquisition measurements, at a single point or for an entire A-line (axial depth), in terms of the displacement and phase in response to tones (or other stimuli). In general, the system noise floor was ~0.8 nm at 0.1 kHz and decreased to below 10 pm from ~10 kHz up to 63 kHz.

For OCT vibrometry, the data acquired by an external trigger provided by the SyncAv software and trigger pulse train was synchronized with the tone stimuli at rates of up to 240 kHz using frequency-doubling software running on a PXI-6221 board (NI, TX, USA). Phase delay in the system was compensated for using custom-built postprocessing software written in MATLAB (R2020a; Natick, MA, USA). The performance of the OCT hardware and VibOCT software was evaluated by measuring the displacement and phase of a small piezoelectric vibrator over a wide range of frequencies and stimulus levels. The OCT displacements and phase were compared to measurements made using a calibrated laser-Doppler vibrometer (LDV; Polytec OFV501/OFV2600, Irvine, CA), and both gave comparable results.

The animal was placed on a two-stage goniometer (07-GON-503, Melles Griot, Carlsbad, CA, USA) that was positioned on top of a 3-axis micro-manipulator (OCT-XYR1, Thorlabs, Germany) mounted on a vibration-isolation table. The head was oriented so that measurements could be made through the intact RWM with the viewing angle adjusted to be close to transverse with respect to the OoC and BM. This viewing angle provided access to BFs in the 40–50 kHz range (see Fig. 1c). Oriented by the real-time OCT images, we chose a region of interest (e.g., BM, OHC-DC-junction, RL) from an A-scan depth profile (Fig. 1d). We then made measurement scans (M-scans), which consist of A-scan depth profiles as functions of time that contain the vibration information. Each B-scan (for imaging) and M-scan (for vibrometry) had 8,192 samples. A second FFT was performed along the time axis of the interferometric phase data for the extraction of vibration displacement and phase (Gao et al., 2013; Lin et al., 2019). The vibration responses to a series of sequential tones were saved to allow further viewing and post-processing at any A-line location. With our custom software, the data could be analyze during the experiment within minutes of being acquired.

### Statistics

To determine the significance of the motion (magnitude and phase) of the each structure (*i.e*., RL_1, 2, 3_, OHC-DC-junction_1, 2, 3_), we performed statistical analysis with multiway (*n*-way) analysis of variance (ANOVA) for testing the effects of multiple factors using the built-in MATLAB function anovan (MATLAB R2020a; Natick, MA, USA). The analysis indicates that the different locations were statistically independent (p < 0.05 criterion). The statistical results are shown in Fig. 5 (RL) and Fig. 7 (OHC-DC-junction).

## Supporting information

Supplemental Information Figurres

## Acknowledgements

We thank John J. Guinan, Jr., for extensive discussion and help in writing the manuscript; Kevin N. O’Connor for the data analysis scripts (SyncAv Toolbox) and editing assistance; Anbuselvan Dharmarajan and Gabriel Alberts for animal surgery; Michael E. Ravicz for technical support; and Andrew A. Tubelli for the OoC-structure drawings (Fig. 1c, g). This work was supported in part by Grant R01 DC07910 from the National Institute on Deafness and Other Communication Disorders (NIDCD) of the NIH, and the Amelia Peabody Charitable Fund.

## Author contributions

N.H.C. and S.P conceived of and designed the project. N.H.C developed the VibOCT software used for these measurements. Both conducted experiments, plotted, and analyzed data. S.P. and N.H.C wrote and edited the manuscript, and S.P. supervised the project.

## Data availability

All data needed to evaluate the conclusions in the paper are present in the paper. Additional data related to this paper may be requested from the authors.

## Additional Information

Supplementary Information is available.

