## Supplemental Information Figurres for "Motion of the Cochlear Reticular Lamina Varies Radially Across Outer-Hair-Cell Rows"

**Supplementary Note 1.** Reticular-lamina (RL) gain measurements at three radial locations, normalized by the high-level basilar-membrane (BM) gain at the junction of the arcuate zone (AZ) and pectinate zone (PZ), $BM_{APJ}$, in three individual animals.

**Supplementary Figures 1–3** present the RL motions at the three radial locations ($RL_3$, $RL_2$, $RL_1$) corresponding to the apical surfaces of the three outer-hair-cell (OHC) rows, from three animals including the one shown in the main manuscript (G637; Fig. 4). The gains and phases from the living (dark colors) and postmortem (PM; faded colors) animals are normalized by the averaged $BM_{APJ}$ gain measured at a high level in order to remove the contribution of the BM traveling wave to RL motion (see Fig. 4).

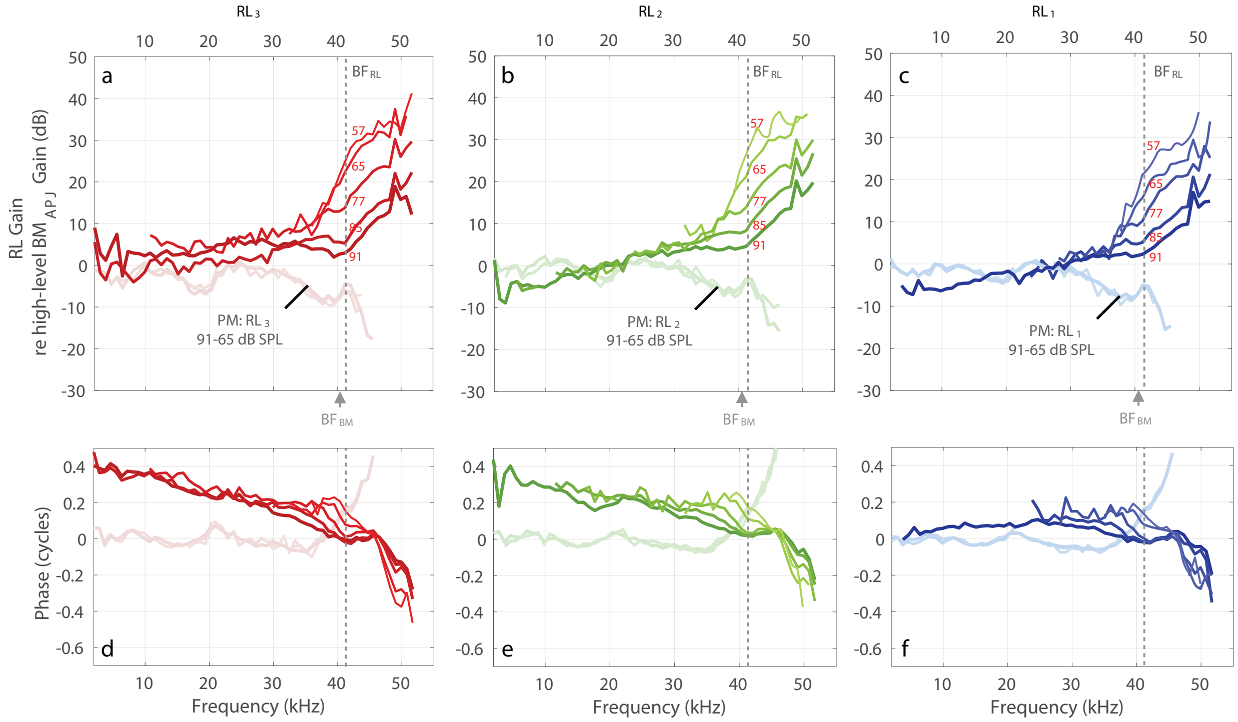


**Supplementary Figure 1.** Specimen G614: **a–c** In vivo and PM gains for $RL_3$, $RL_2$, and $RL_1$, respectively, normalized by the high-level $BM_{APJ}$ gain (in dB). **d–f** The phase responses corresponding to panels **a–c**. Note that the baseline high-level $BM_{APJ}$ gain used for normalization was calculated as the average of five measurements made at 91 dB SPL. All frequency axes in this figure are on a linear scale.

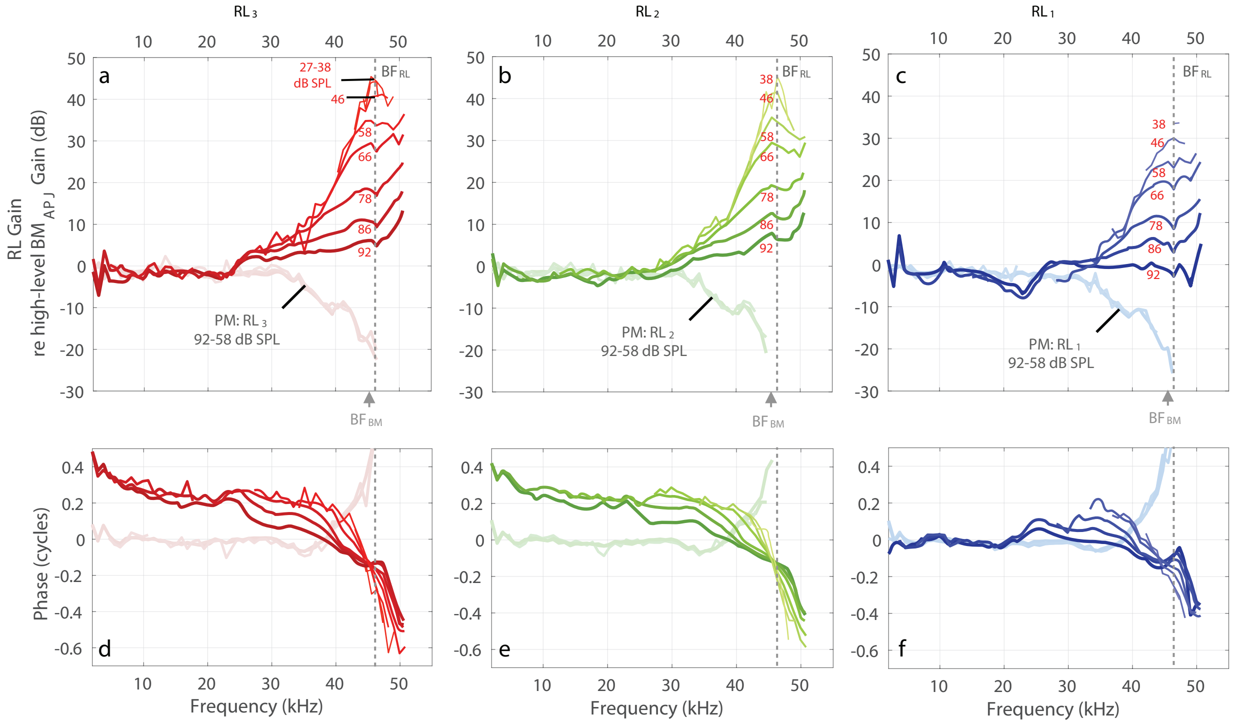


**Supplementary Figure 2.** Specimen G637: **a–c** In vivo and PM gains for $RL_3$, $RL_2$, and $RL_1$, respectively, normalized by the high-level $BM_{APJ}$ gain (in dB). **d–f** The phase responses corresponding to panels **a–c**. Note that the baseline high-level $BM_{APJ}$ gain used for normalization was calculated as the average of ten measurements made at 92 dB SPL. All frequency axes in this figure are on a linear scale. This dataset is also shown in Fig. 4 of the main manuscript.

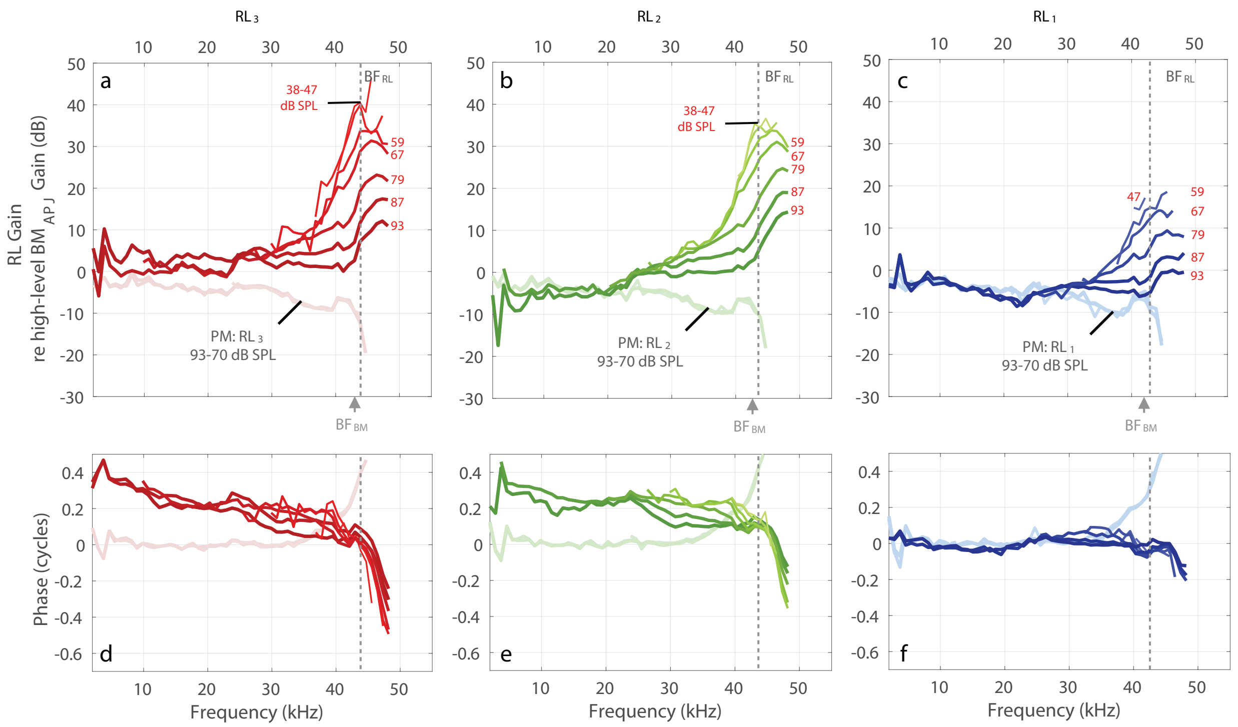


**Supplementary Figure 3.** Specimen G645: **a–c** In vivo and PM gains for $RL_3$, $RL_2$, and $RL_1$, respectively, normalized by the high-level $BM_{APJ}$ gain (in dB). **d–f** The phase responses corresponding to panels **a–c**. Note that the baseline high-level $BM_{APJ}$ gain used for normalization was calculated as the average of four measurements made at 93 dB SPL. All frequency axes in this figure are on a linear scale.

**Supplementary Note 2.** Gain measurements at three transverse locations in the organ of Corti (OoC), normalized by sound pressure, in four individual animals.

**Supplementary Figures 4–7** present the motions of three different transverse locations: $RL_3$, the junction between OHC row and its connected Deiters' cell (OHC-DC-junction), and $BM_{APJ}$, in four animals. These figures show similar results to Fig. 10 in the main manuscript.

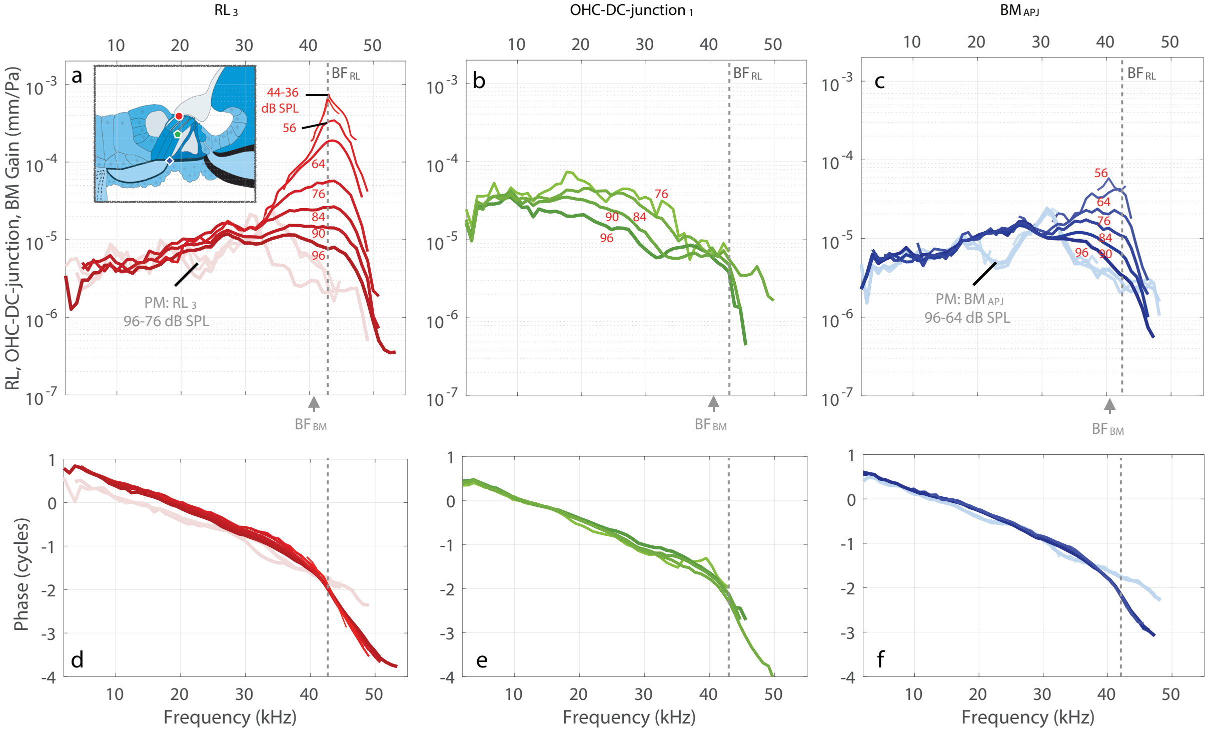


**Supplementary Figure 4.** Specimen G612: **a–c** In vivo (darker colors) and PM (faded colors) gains are shown for $RL_3$ (red), OHC-DC-junction$_1$ (green), and $BM_{APJ}$ (blue), all relative to the sound pressure (in units of mm/Pa). The inset drawing in panel **a** shows the measurement locations for $RL_3$ (red circle), OHC-DC-junction$_1$ (green pentagon), and $BM_{APJ}$ (blue diamond). **d–f** The respective phase responses corresponding to **a–c**. Note that the available stimulus levels vary across the structures due to different signal-to-noise ratios. No PM OHC-DC-junction$_1$ measurements were available for this specimen (**b** and **e**).

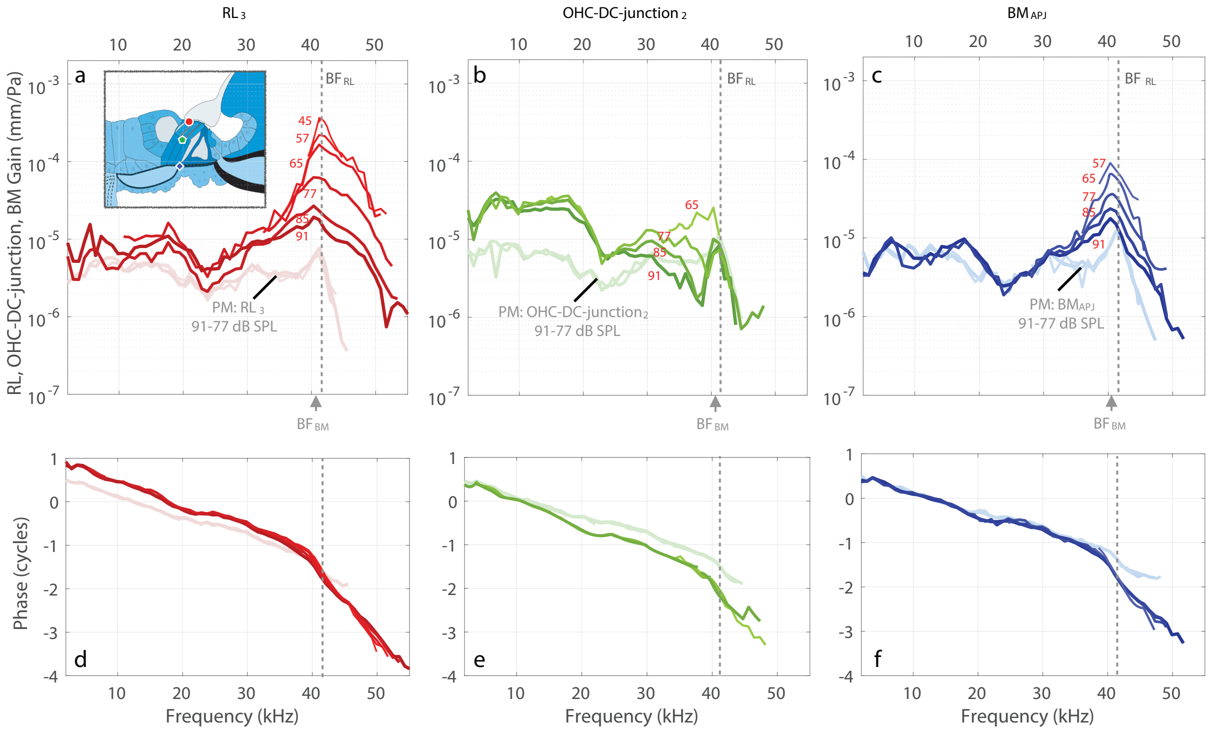


**Supplementary Figure 5.** Specimen G614: **a–c** In vivo (darker colors) and PM (faded colors) gains are shown for $RL_3$ (red), OHC-DC-junction$_1$ (green), and $BM_{APJ}$ (blue), all relative to the sound pressure (in units of mm/Pa). The inset drawing in panel **a** shows the measurement locations for $RL_3$ (red circle), OHC-DC-junction$_1$ (green pentagon), and $BM_{APJ}$ (blue diamond). **d–f** The respective phase responses corresponding to **a–c**. Note that the available stimulus levels vary across the structures due to different signal-to-noise ratios.

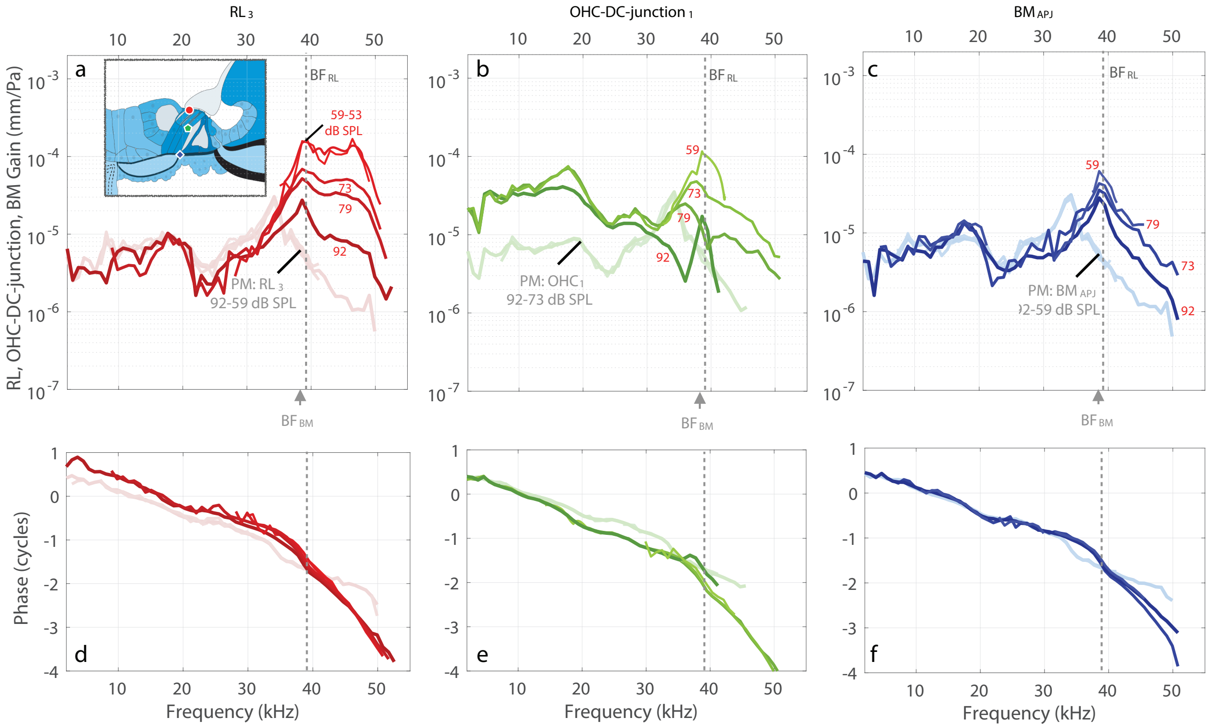


**Supplementary Figure 6.** Specimen G619: **a–c** In vivo (darker colors) and PM (faded colors) gains are shown for $RL_3$ (red), OHC-DC-junction$_1$ (green), and $BM_{APJ}$ (blue), all relative to the sound pressure (in units of mm/Pa). The inset drawing in panel **a** shows the measurement locations for $RL_3$ (red circle), OHC-DC-junction$_1$ (green pentagon), and $BM_{APJ}$ (blue diamond). **d–f** The respective phase responses corresponding to **a–c**. Note that in this animal the $RL_3$ results show broad tuning curves in panel **a**.

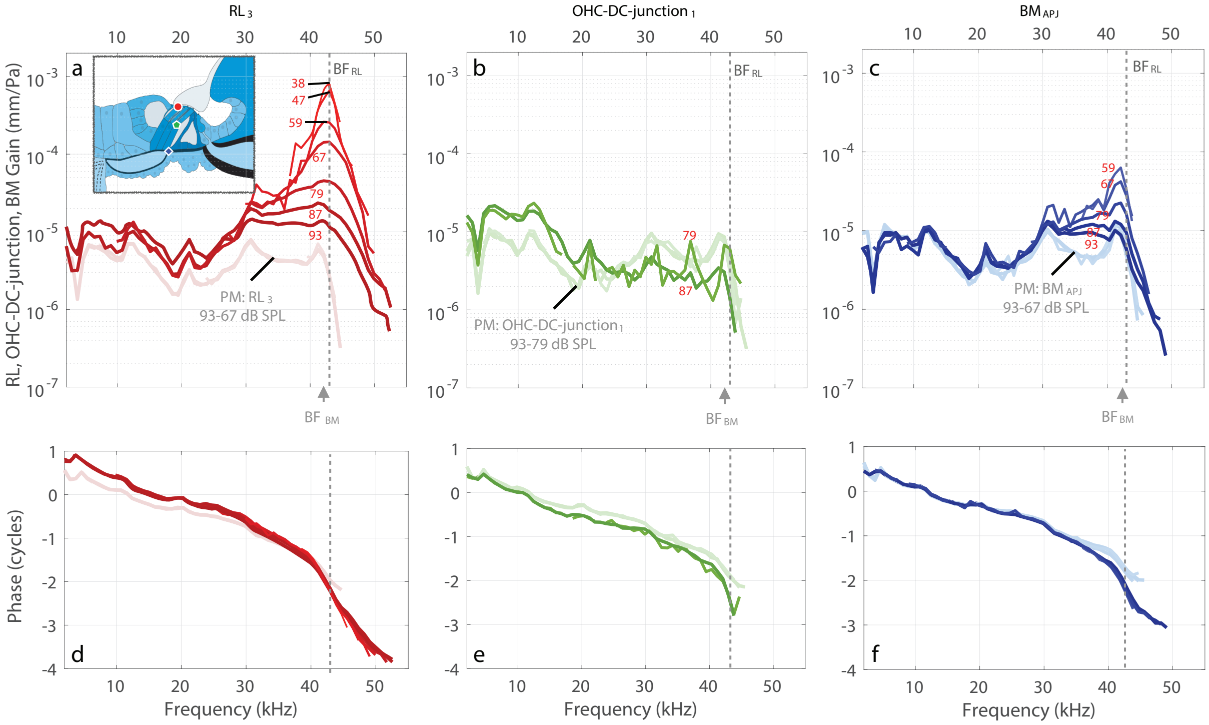


**Supplementary Figure 7.** Specimen G645: **a–c** In vivo (darker colors) and PM (faded colors) gains are shown for $RL_3$ (red), $\text{OHC-DC-junction}_1$ (green), and $BM_{APJ}$ (blue), all relative to the sound pressure (in units of mm/Pa). The inset drawing in panel **a** shows the measurement locations for $RL_3$ (red circle), $\text{OHC-DC-junction}_1$ (green pentagon), and $BM_{APJ}$ (blue diamond). **d–f** The respective phase responses corresponding to **a–c**. Note that the available stimulus levels vary across the structures due to different signal-to-noise ratios.

**Supplementary Note 3.** BM gain measurements at three radial locations, normalized by the high-level $BM_{APJ}$ gain, in specimen G637.

**Supplementary Figure 8** presents the motions at three radial points along the BM ($BM_{PZ}$, $BM_{APJ}$, and $BM_{AZ}$), normalized by the high-level $BM_{APJ}$ gain (in dB), in animal G637. For all three BM locations, the group delay (i.e., the negative slope of the phase-versus-frequency function in Supplementary Figure 8d–f) became smaller as the sound level increased, but the group delays for the BM were about two times smaller than those of the RL in Fig. 4f–h of the main manuscript. The BM group delays ~0.06 cycles and RL group delays ~0.14 cycles at the frequency 35 kHz between 92 and 66 dB SPL stimulus level which could be include propagation delay (timing difference) from the BM to RL structure.

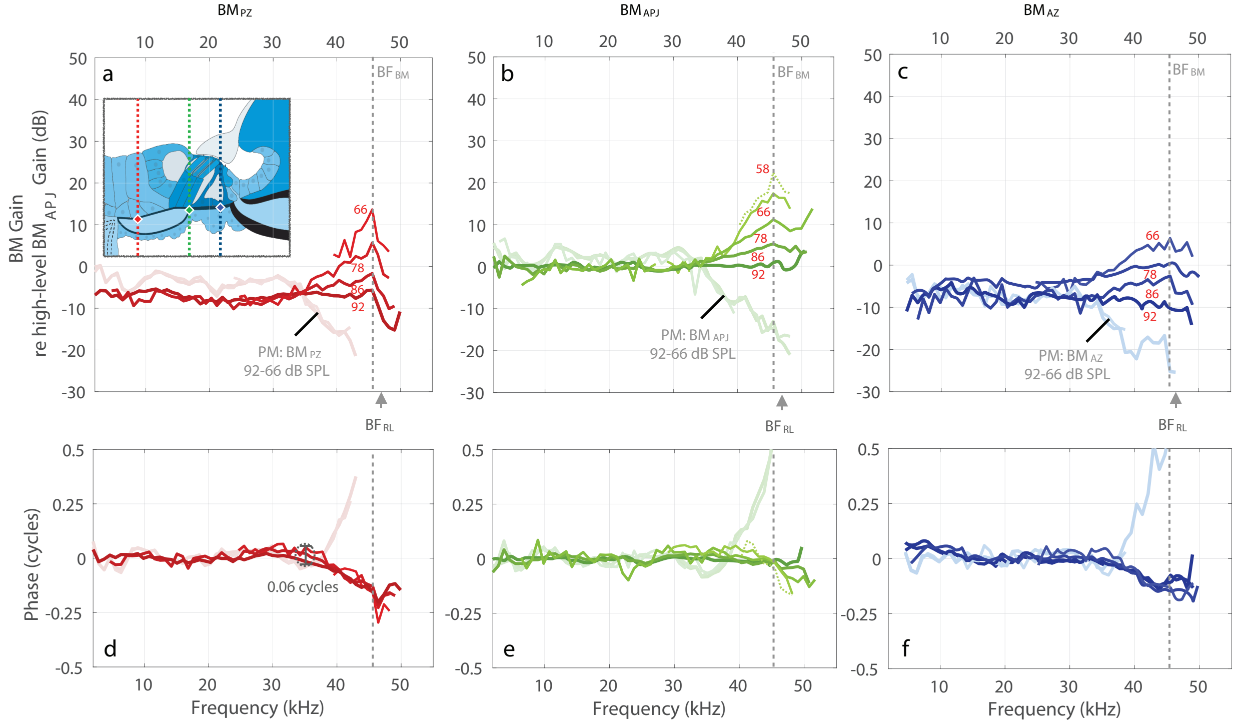


**Supplementary Figure 8.** Specimen G637: **a–c** In vivo and PM gains for $BM_{PZ}$, $BM_{APJ}$, and $BM_{AZ}$, respectively, normalized by the high-level $BM_{APJ}$ gain (in dB). **d–f** The phase responses corresponding to panels **a–c**. Note that the baseline high-level $BM_{APJ}$ gain used for normalization was calculated as the average of ten measurements made at 92 dB SPL. All frequency axes in this figure are on a linear scale.
